# EEG-based neurofeedback with network components extraction: a data-driven approach by multilayer ICA extension and simultaneous EEG-fMRI measurements

**DOI:** 10.1101/2021.06.20.449196

**Authors:** Takeshi Ogawa, Hiroki Moriya, Nobuo Hiroe, Motoaki Kawanabe, Jun-ichiro Hirayama

## Abstract

Several studies have reported advanced treatments for depressive symptoms, such as real-time neurofeedback (NF) with functional MRI (fMRI) and/or electroencephalogram (EEG). NF focusing on a regularization of brain activity associated with the amygdala or functional connectivity (FC) between the executive control network (ECN) and default mode network (DMN) has been applied to reduce depressive symptoms. However, it is practically difficult to install the fMRI-NF system and to consistently provide treatment, because of high cost. Additionally, no practical signal processing techniques have been developed extracting FC-related features from EEG signals, particularly when no physical forward models are available. In this regard, stacked pooling and linear components estimation (SPLICE), recently proposed as a multilayer extension of independent component analysis (ICA) and related independent subspace analysis (ISA), can be a promising alternative. The resting-state EEG network features can be correlated with fMRI network activity corresponding to the DMN or ECN. This may enable the modulation of the target FC-related features in EEG-based NF.

In this study, we developed a real-time EEG NF system for improving depressive symptoms by using the SPLICE. Utilizing information from the fMRI biomarkers, we evaluated our paradigm for effectiveness with regard to upregulation of the dorsolateral prefrontal cortex /middle frontal gyrus or downregulation of the precuneus/posterior cingulate cortex. We conducted an NF experiment in participants with subclinical depression; the participants were divided into the NF group (n=8) and the sham group (n=9). We found a significant reduction and a large effect size in the rumination response scale (RRS) score (reflection) in the NF group, compared to the sham group.

However, we did not find a significant relationship between the training score and difference in symptoms. This suggests that increased controllability of the EEG signals did not directly reduce the RRS reflection score. This could be due to various reasons such as improper feature extraction, individual differences, and the targeted brain regions. In this paper, we also discuss the possible ways to modify our NF protocol including the design of the experiment, sample size, and online processing. We then discuss way to improve the NF training, based on our results.

## 1. Introduction

Advanced treatments for depressive symptoms have been formulated using brain or physiological measurements instead of traditional medical treatments. One of the noninvasive ways to accurately monitor brain activity that may represent depressive symptoms is functional MRI (fMRI). This technique is well suited because of its high spatial resolution, even when applied to deep brain areas. Recently, several groups have proposed real-time fMRI neurofeedback (NF) paradigms to regulate participants’ brain signals for converting depressive brain states into healthier states (Young et al., 2017; Yamada et al., 2017; Taylor et al., 2021; Takamura et al., 2020). It is assumed that at the end of the NF training period, improvement in the controllability of participant brain activity could help reduce their depressive symptoms.

The fMRI NF, focusing on regularizing functional connectivity (FC) between the executive control network (ECN) and default mode network (DMN), has been applied to alleviate depression (Yamada et al., 2017; Taylor et al., 2021; Tsuchiyagaito et al., 2021). The relationship between the time courses of the ECN and DMN in healthy individuals is generally negatively correlated, and is associated with the distinction between internal (own thoughts) and external (perceptions of the external environment) contents (Northoff, 2016). In contrast to healthy individuals, patients with depressive disorders tend to be dysbalanced, such as positively or not significantly correlated, and show increased self-focus (rumination, guilt, fear) and decreased environment-focus (social withdrawal, psychomotor retardation) (Drysdale et al., 2017). This relationship was elucidated using an fMRI biomarker comprised of FCs; utilizing data-driven approaches, this biomarker was used to successfully distinguish between patients with melancholic depression and healthy controls (Ichikawa et al., 2020; Yamashita et al., 2020). They found the FCs including the dorsolateral prefrontal cortex (DLPFC)/middle frontal gyrus (MFG) in ECN and the precuneus/posterior cingulate cortex (PCC) in DMN. According to the results of the fMRI biomarker analysis, NF with a specific FC, termed as FC neurofeedback (FCNef), shows potential to be a novel treatment modality for depression, and can be applied in a targeted manner.

However, real-time fMRI NF takes high costs; it is practically difficult to install this system and constantly provide treatment. This is a significant obstacle for applying this technique to treat depressive symptoms. On the other hand, real-time NF using electroencephalography (EEG) has been proposed as an alternative approach, such as NF applied with frontal alpha asymmetry (Wang et al., 2019), peak alpha frequency (Yu et al., 2020), and alpha-theta ratio (Cheon et al., 2016). In particular, frontal alpha asymmetry EEG NF has been shown to modulate left-side alpha power associated with brain activity in the left amygdala (Davidson et al., 1990). EEG has many practical benefits such as reasonable cost, high temporal resolution, and fewer physical constraints; its main limitation is low-spatial resolution. Recent studies of EEG-fMRI data have reported the possible effectiveness of real-time EEG-fMRI NF paradigms for depression (Zotev et al., 2014; Zotev et al., 2016; Zotev et al., 2020) and improved reliability of EEG signatures for NF. In these NF training studies, participants were asked to imagine a positive autographic memory, such as a happy event, and were instructed to increase brain activity in the left amygdala and simultaneously try to increase frontal alpha asymmetry. This NF modality aims to reinstate healthy brain function in the left amygdala.

However, in contrast to real-time fMRI NF or FCNef, an EEG NF using network features has been poorly investigated. This is mainly because there are no practical signal processing techniques extracting FC-related features from EEG, particularly when no physical forward models are available. Especially in real-time process, a brain signal source localization is a difficult issue that requires to solve inverse problems with constraints. In fact, most studies have used sensor-level features such as time-varying powers of particular frequency bands, or have sought to extract source-level features by a blind source separation (BSS) technique such as independent component analysis (ICA). Neither method is designed to detect EEG features that reflect FC and modular organization related to resting-state networks. Therefore, it is imperative to develop a new BSS-based technique combined with a signal model that explicitly assumes network structures at the source level related to FC. In this regard, stacked pooling and linear components estimation (SPLICE; Hirayama et al., 2017), which has been recently proposed as a multilayer extension of ICA and related independent subspace analysis (ISA; Hyvärinen & Hoyer, 2000), can be a promising alternative. However, its application to resting-state EEG has not yet been investigated. We expected that if the extracted EEG network features correlate with fMRI network activity corresponding to the DMN or ECN, these features may be used to modulate the target networks in EEG-based NF training.

In this study, we attempted to apply an EEG network estimated by SPLICE for a real-time EEG NF paradigm. We have developed EEG-based NF training to improve depressive symptoms in healthy individuals with subclinical depression. We evaluated the effectiveness of EEG NF in modulating estimated brain activity; targeted modulation included upregulation of DLPFC/MFG activity or the downregulation of precuneus/PCC activity, which was expected to reduce depressive symptoms. We conducted a four-day EEG-NF study with two groups (NF, sham) of participants with sublicinal depression; we targeted specific regions (DLPFC/MFG or precuneus/PCC) while applying an FCNef paradigm (Yamada et al., 2017; Taylor et al., 2021). To train the SPLICE model, participants completed a two-day EEG-fMRI experiment; in this experiment, we obtained simultaneously recorded EEG-fMRI data at rest and during a working memory task such as the N-back task which evoked brain activity in the ECN. We selected EEG network features that correlated with fMRI signals in the DLPFC or precuneus/PCC for NF training, by modeling SPLICE from the resting-state data. Scores on the Beck Depression Inventory-II (BDI-II; Beck et al., 1996) and the Rumination Response Scale (RRS; Hasegawa, 2013; Treynor et al., 2003) were measured pre-and post-NF training. Finally, we evaluated the effect size of the difference between pre-and post-training among NF and sham groups. In this paper, we present the results of these experiments and also discuss the possibility of formulating an EEG NF protocol using EEG network features relevant to fMRI network information.

## 2 Materials and Methods

### 2.1 SPLICE

SPLICE (Hirayama et al., 2017) is a multilayer extension of ICA that is based on a probabilistic model that simplifies the generative process of EEG. This concept can be mathematically described as follows:

Let x and s denote the vectors of the EEG measurements and unobserved source signals, respectively, at a single time point. Then, we assume a linear model (first layer) similar to ICA, given by

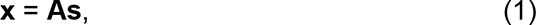

where the mixing matrix A is a square matrix and invertible; the inverse **W**: = **A**^-1^ is called the demixing matrix. Both **x** and **s** are zero-mean, without loss of generality, by subtracting the sample mean in advance from the original data vectors. Note that the sources (entries in **s**) lack explicit physical mapping to the cortex. However, they can still be interpreted as reflecting cortical activations based on their associated topographies (i.e., corresponding columns of **A**), exactly as in ICA. For mathematical convenience, the number of sources, d, is assumed to be the same as the dimensionality of **x**; in practice, we apply PCA beforehand to determine an appropriate number of sources as customarily done in ICA.

Additional layers are introduced on top of the first layer to model the statistical dependency among the sources and the parametric structures behind it. Specifically, SPLICE introduces 1) the functional modularity of sources and 2) the intrinsic correlations (coactivations) among modules into the multilayer generative model. Both are reasonable assumptions, particularly for resting-state EEG data. This is because cortical activity at rest is known to exhibit functional organization with multiple modules (e.g., resting-state network) that are often mutually correlated or anticorrelated.

First, we divide the d first-layer sources into m modules **s**[*j*] without overlapping, where **s**[*j*] denotes the vector consisting of *dj* sources belonging to module j 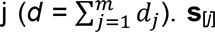 represents a *dj*-dimensional subspace spanned by the corresponding columns in **A**.

Then, we assume that the (squared) L2-norms ||**s**[*j*]||^2^ of modules **s**[*j*] are generated by an additional ICA-like linear generative model with pointwise nonlinearity, such that generated by an additional ICA-like linear generative model with pointwise nonlinearity, such that

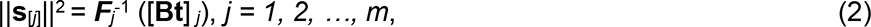

where an appropriate link function ***F****j* is used for a convention between non-negative (norm) and real random variables (we set ***F****j* = log in our analysis); **B** and **t** constitute the invertible mixing matrix, and source vector (and **V**:= **B**^-1^ is a demixing matrix) of this layer, and [·] *j* denotes the *j*-th entry of a vector. Then, the norms are either mutually uncorrelated or correlated, depending on specific values of **B** (see below). Here, the squared L2-norm of a vector is simply the sum-of-squares of entries and it represents the total power of the sources within each module. In this study, we focus on modeling and analyzing the module norms (powers), making a minimal assumption on module direction **s**[*j*]/||**s**[*j*]||. Thus, we assume module directions **s**[*j*]/||**s**[*j*]|| are mutually independent and each follows a spherically uniform distribution, which maximizes the entropy under restriction of unit norm constraints. This is equivalent that we assume every modules vectors **s**[*j*] to be spherically distributed. Note that the model can be recursively extended to have additional layers to further model the dependency in higher-layer sources (Hirayama et al., 2017). However, we assume Eq.(2) describes the top layer in the present study. Thus, the entries in t are assumed to be independent of each other, given a typical prior in ICA (e.g., the one corresponding to the standard tanh nonlinearity).

Figure 1. The model consists of three layers, as follows: two linear layers, and an intermediate L2-pooling layer for calculating the norms. Here, we define **y**: = **Bt**. If the highest layer **B** is not diagonal, the modules are mutually independent, and the model reduces to independent subspace analysis (Hyvärinen & Hoyer, 2000). If not diagonal, the modules are not independent, and are thus expected to be suitable for representing correlated or anticorrelated cortical modules. Note that modeling non-diagonal **B** affects the higher layer and influences the first layer through the joint learning of the two layers (see below) because it makes a different (possibly improved) prior assumption on the first-layer sources from that of ICA or ISA.

**Figure 1.**
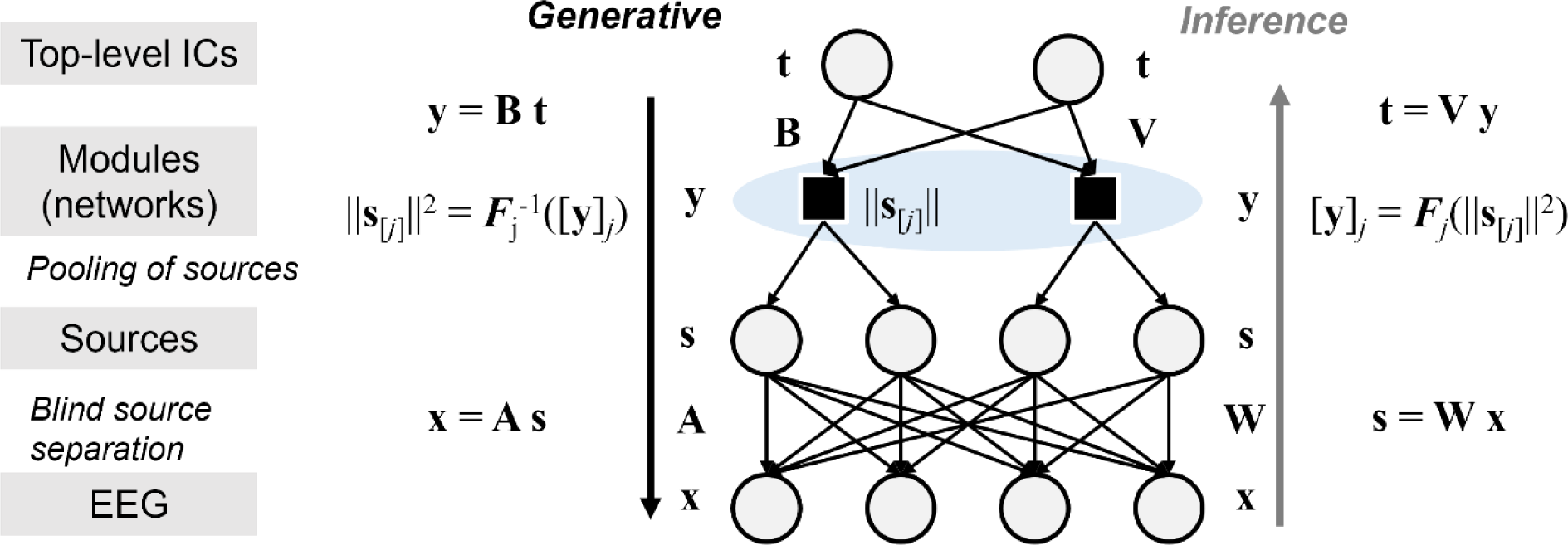
Structure of stacked pooling and linear components estimation (SPLICE). **s**, unobserved source signal; **A**, mixing matrix; **W**, demixing matrix of **A**; **x**, EEG measurements; **B**, invertible mixing matrix; **t**, source vector; ***F****j*, link function; **V**, demixing matrix of **B**.

A striking property of SPLICE is that both parameter estimation (learning) and posterior inference on latent variables can be performed in a principled manner without resorting to any approximative techniques or heavy numerical computation. Learning is done based on conventional maximum likelihood (ML) estimation. The corresponding loss function, i.e., the negative log-likelihood for a single datum *L*(**x**): = − *ln p*(**x**) + const. where *p*(**x**) is the probability density function of **x**, is analytically given by

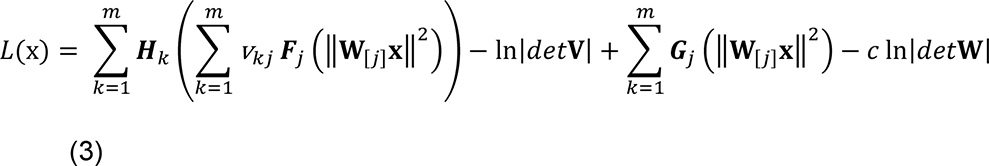

where ***H*** and ***G*** are functions derived from the prior distribution of ***t*** and the link function ***F****j*. The dimensionality of every module (subspace) *dj* is adaptively determined before ML estimation using a clustering-like technique (see Hirayama et al., 2017 for details).

Once demixing matrices **W** and **V** are learned, the generative process can be readily inverted by computing **s** = **Wx** and **y** = **Bt**, with the L2-pooling ||**s**[*j*]||^2^ (followed by ***F****j*) between them. In practice, the pooled outputs (squared norm) may fluctuate too heavily owing to the noisy nature of the EEG technique. Thus, as a slight extension of the original SPLICE, we also implemented the L2-pooling across successive time points for smoothing; this was done in a principled manner by modifying the model and likelihood, so that the top linear layer determines the L2 norm of **s**[*j*] that now concatenates several instances of the *dj* sources across time points. As noted above, pooling just means taking the sum of squares (of the filtered outputs). Thus, the squared module norm essentially provides the total power or energy of a source module’s activity. Therefore, we focused primarily on the estimated module norms as the main output of the analysis, rather than the top-layer sources **t** which the multilayer computation eventually produces, because we expected to associate EEG features with the activity of cortical modules.

### 2.2 Experiment procedures

#### 2.2.1 Participants

The workflow pertaining to this experimental procedure is illustrated in Figure 2A. The experiment consisted of three steps, as follows: “Screening,” “EEG-fMRI experiment,” and “NF training experiment.” All participants were recruited by screening questionnaires and clinician assessment (n = 102; 49 women, 53 men; mean age = 29.88 ± 9.8 years) at the Screening step. Before the EEG-fMRI experiment, participants, who (a) were prone to suicidal thoughts as measured in the BDI, (b) had current/past mental/psychiatric diseases, (c) did not understand the Japanese language, (d) showed a BDI score less than 3, or (e) declined to attend due to time unavailability or other personal reasons, were excluded. Twenty-eight participants with subclinical depression, designated as “subclinical individuals”, were included in the EEG-fMRI experiment which was conducted for two days. Furthermore, we excluded participants who had excessive body/head movements in fMRI scanning and declined to attend due to time unavailability or other personal reasons. Finally, 17participants (NF group, n = 8; sham group: n = 9) completed the NF training experiment conducted for four days. All participants provided written informed consent prior to participating in the experiment. After each experiment, all participants received cash remuneration for their involvement. This study was approved by the ethics committee of the Advanced Telecommunications Research Institute International (ATR) and was conducted in accordance with the Declaration of Helsinki.

**Figure 2.**
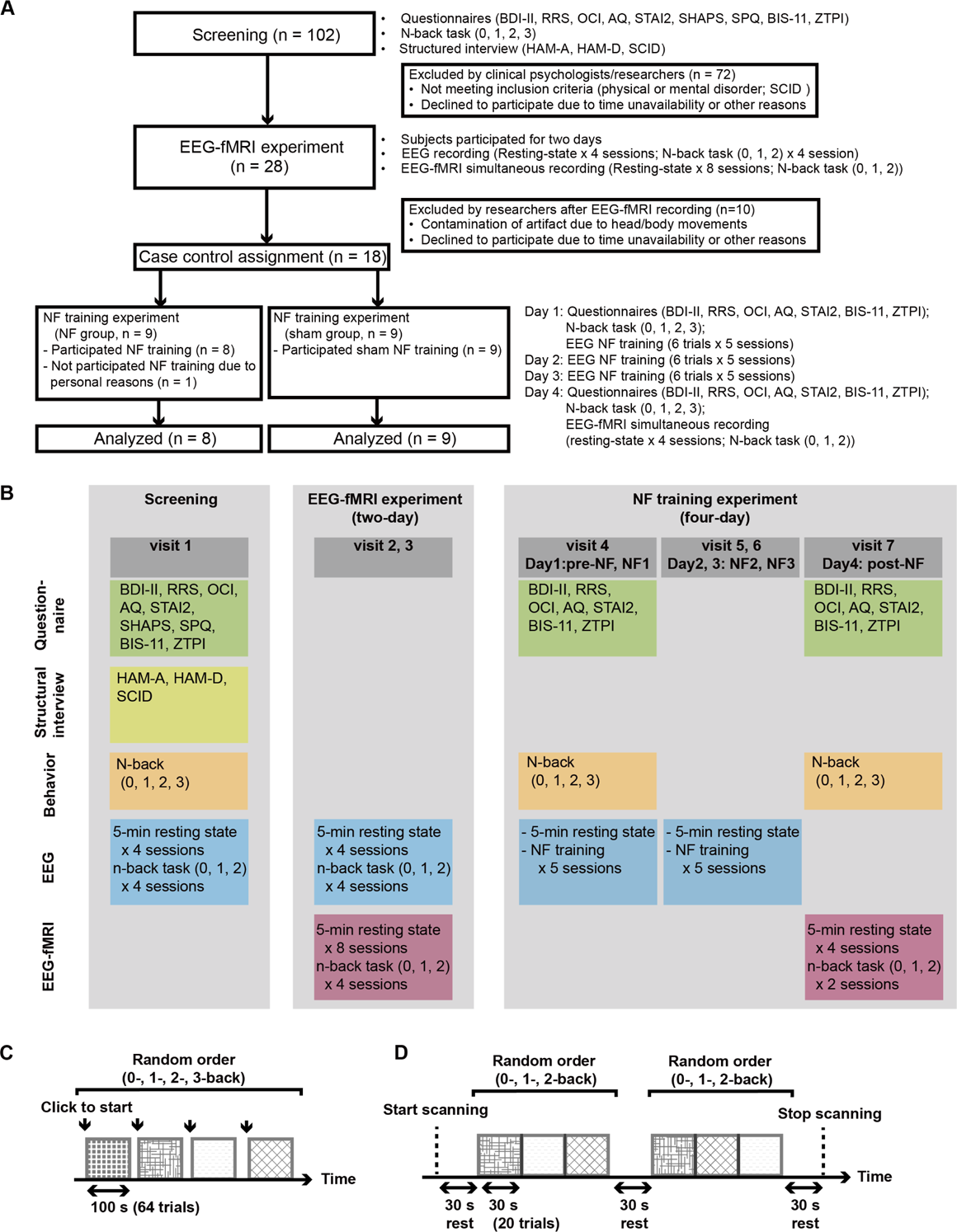
Experimental workflow. (**A**) A schema of the experimental procedure (**B**) A summary of measurements made at each visit (**C**) A diagram depicting the N-back task for the behavioral experiment at visits 1, 4 and 7. (**D**) A diagram depicting the N-back task for electroencephalography (EEG)/EEG-functional magnetic resonance imaging (fMRI) experiment at visits 2, 3, and 7. BDI-II: Beck Depression Inventory II; RRS: Rumination Response Scale; OCI: Obsessive-Compulsive Inventory; AQ: Autism-Spectrum Quotient; STAI2: State-Trait Anxiety Inventory, SHAPS: Snaith-Hamilton Pleasure Scale; SPQ: Schizotypal Personality Questionnaire; BIS-11: Barratt Impulsiveness Scale 11; ZTPI: Zimbardo Time Perspective Inventory; HAM-D: 21-item Hamilton Depression Rating Scale; HAM- A: Hamilton Anxiety Rating Scale; SCID: Structured Clinical Interview for DSM-IV-TR Axis disorders (SCID); NF: Neurofeedback.

#### 2.2.2 Psychological questionnaires

In the Screening experiment (visit 1), all participants completed self-reported questionnaires that included queries pertaining to demographic characteristics, BDI, RRS, Obsessive-Compulsive Inventory (OCI), Autism-Spectrum Quotient (AQ), State-Trait Anxiety Inventory (STAI2), Snaith-Hamilton Pleasure Scale (SHAPS), Schizotypal Personality Questionnaire (SPQ), Barratt Impulsiveness Scale 11 (BIS-11), Zimbardo Time Perspective Inventory (ZTPI); participants were rated using the 21-item Hamilton Depression Rating Scale (HAM-D), the Hamilton Anxiety Rating Scale (HAM-A), and the Structured Clinical Interview for DSM-IV-TR Axis disorders (SCID). To evaluate the effects of the EEG NF training, participants completed the BDI, RRS, OCI, AQ, STAI2, BIS-11, and ZTPI on Day 1 (visit 4) and Day 4 (visit 7) of the NF training experiment (Figure 2B). We found a significant positive correlation between the BDI and RRS score (total: 0.66; brooding: 0.44; reflection: 0.59), including the subscale, which presented in Supplementary Figure 1.

#### 2.2.3 N-back task

The N-back task is a cognitive task associated with the functioning of the working memory. We conducted the N-back task test to induce brain activities with anti-correlation between the ECN and DMN (Yamada et al., 2017; Liu et al., 2017; Cao et al., 2021). This enabled us to select a SPLICE module for the NF training. Participants completed a behavioral experiment, such as N-back tasks, outside the MRI screener during the Screening experiment (visit 1) (baseline), and during the NF training experiment (visit 4: Day 1, visit 7: Day 4). We used these data to evaluate changes in cognitive performance resulting from the NF training (see Figure 2C). Customized code generated using Presentation ® software (Version 18.0, Neurobehavioral Systems, Inc., Berkeley, CA) controlled the N-back task and to display the task on the monitor. When ready to start the N-back task, participants clicked the left button of a mouse to begin each session, shown in Figure 2C. The participants were instructed to relax and simply focus at the center of the monitor. In the 0-back condition, participants were asked to promptly click the left mouse button when the number “0” appeared on the monitor. In the 1-back, 2-back, and 3-back conditions, participants were asked to click the left mouse button promptly when matching a currently displayed number and a number presented one, two, or three trials prior. A single session consisted of four blocks with four randomly aligned conditions (0-, 1-, 2-, and 3-back). A single trial had an inter-trial interval of 1300 ms duration, after which a number from 1 to 9 was displayed at the center of the monitor for 200 ms. In total, a single block had 64 trials that lasted for 100 s; therefore, it took approximately 13 min to complete two sessions of this task.

In the EEG-fMRI experiment, participants also completed the N-back task with EEG recording outside the MRI scanner and fMRI scanning inside the MRI scanner (see Figure 2D). When in the MRI scanner, participants were asked to press the response buttons using an MRI-compatible response device (HHSC-2 × 2, Current Designs, Inc., PA, USA). A single session consisted of the following: a first 30-s resting period (a white fixation cross at the center of the screen), three randomly ordered blocks of three conditions (0-, 1-, and 2-back), a second 30-s resting period, three blocks of N-back tasks, and a third 30-s resting period. At the beginning of each block, participants were informed of a rule corresponding to the condition, and were then asked to complete 20 trials. It took participants 5 min 45 s to complete a single session.

### 2.3 Data acquisitions

Figure 2B summarizes the measurements performed at each visit. MRI data were acquired with a 3T MRI scanner, MAGNETOM Verio (Siemens, Erlangen, Germany), using a Siemens 12-channel head coil installed at the Brain Activity Imaging Center located in the ATR. High-resolution, T1-weighted structural images (TR = 2250 ms, TE = 3.06 ms, flip angle = 9 °, inversion time = 900 ms, matrix = 256 × 256, 208 sagittal slices, 1 mm isotropic) were acquired for normalization to a standard brain for echo-planar image (EPI) registration. Functional images such as those using fMRI data were acquired with an EPI sequence (TR = 2450 ms, TE = 30 ms, flip angle = 80 °, matrix = 64 × 64, field of view = 192 mm, slice thickness = 3.2 mm, 35 axial slices, scan sequence: ascending). We measured 150 volumes (6 min 7 s) for the resting state and 141 volumes (5 min 45 s) for the N-back task.

EEG signals were measured using an MR-compatible amplifier (BrainAMP MR plus, Brain Products GmbH, Germany) and a 64-channel EEG electrode cap (BrainCap MR, Brain Products GmbH, Germany); this included 63 EEG channels and one electrocardiogram (ECG) channel. The EEG cap was correctly placed on the participant’s head as per the International 10-10 system. The ground electrode and online reference electrode were placed at AFz and FCz positions, respectively. For safety reasons, the impedance of all the EEG electrodes and the ECG electrode were maintained below 10 kΩ and 50 kΩ, respectively, throughout the experiment. We placed the ECG electrode on the participant’s back to enable the subsequent correction of ballistocardiographic artifacts. Using the Brain Vision Recorder (Brain Products GmbH, Germany), raw data were recorded with a 5 kHz sampling frequency, with filtering between 0.1 and 250 Hz. The amplifier system was placed on the MRI scanner beside the subject’s head during fMRI scanning with fixation of MR-compatible sandbags to reduce artifacts caused by the tinny vibration of the scanner. The EEG recording system clock was synchronized using a SyncBox device (Brain Products GmbH, Germany) and the MRI scanner’s 10 MHz master synthesizer to achieve phase synchronization clocks for digital sampling between the MRI data and the EEG system. To remove artifacts caused by fMRI scanning from the EEG data, we simultaneously recorded each TR trigger signal to mark the onset of every instance of single fMRI volume acquisition. During the EEG-fMRI data acquisition, we used a projector (DLA- X7-B, JVC; frame rate = 60 Hz) to present the visual stimulation on an opaque screen.

The EEG-fMRI experiment was conducted as follows: first, EEG measurements of the resting-state (two sessions) and the N-back task (0-, 1-, 2-back; two sessions) were obtained for each participant in a soundproof shield room. A single session of the resting-state and that of the N-back task took five minutes and six minutes, respectively. Second, participants moved to the MRI scanner and completed the resting-state (two sessions), the N-back task (two sessions), the T1-weighted structural image scan, and the resting state (two sessions). Finally, EEG data (resting-state: four sessions; N-back task: four sessions) and EEG-fMRI data (resting-state: eight sessions; N-back task: four sessions) were obtained and used to estimate the SPLICE filter for the NF.

During the NF training experiment, EEG signals were measured in the soundproof shield room from Day 1 to Day 3; EEG-fMRI data were obtained in the MRI scanner on Day 4 as well as during the EEG-fMRI experiment. The resting-state EEG signals were measured for five minutes during the NF training (Days 1, 2, and 3). Subsequently, the NF training was conducted for each participant for nine minutes in a single session; each participant completed five sessions.

### 2.4 Data analysis

#### 2.4.1 Preprocessing of fMRI data

Figure 3A summarizes the workflow of preprocessing of the EEG and fMRI data and the integration process. The preprocessing parameters and methods were as per previous studies (Ogawa et al., 2018; Aihara et al., 2020). The data were processed using SPM8 toolbox in MATLAB software (Wellcome Trust Centre for Neuroimaging). The first four volumes were removed to allow for T1 equilibration. The remaining data were subjected to slice-timing correction and realignment of the mean image of that sequence to compensate for head motion. The structural image was coregistered to the mean functional image and segmented into three tissue classes in the MNI space. The functional images were normalized and single voxels were resampled with a 2 × 2 × 2 mm grid. Finally, the normalized images were spatially smoothed using an isotropic Gaussian kernel of 8 mm full-width at half maximum.

**Figure 3.**
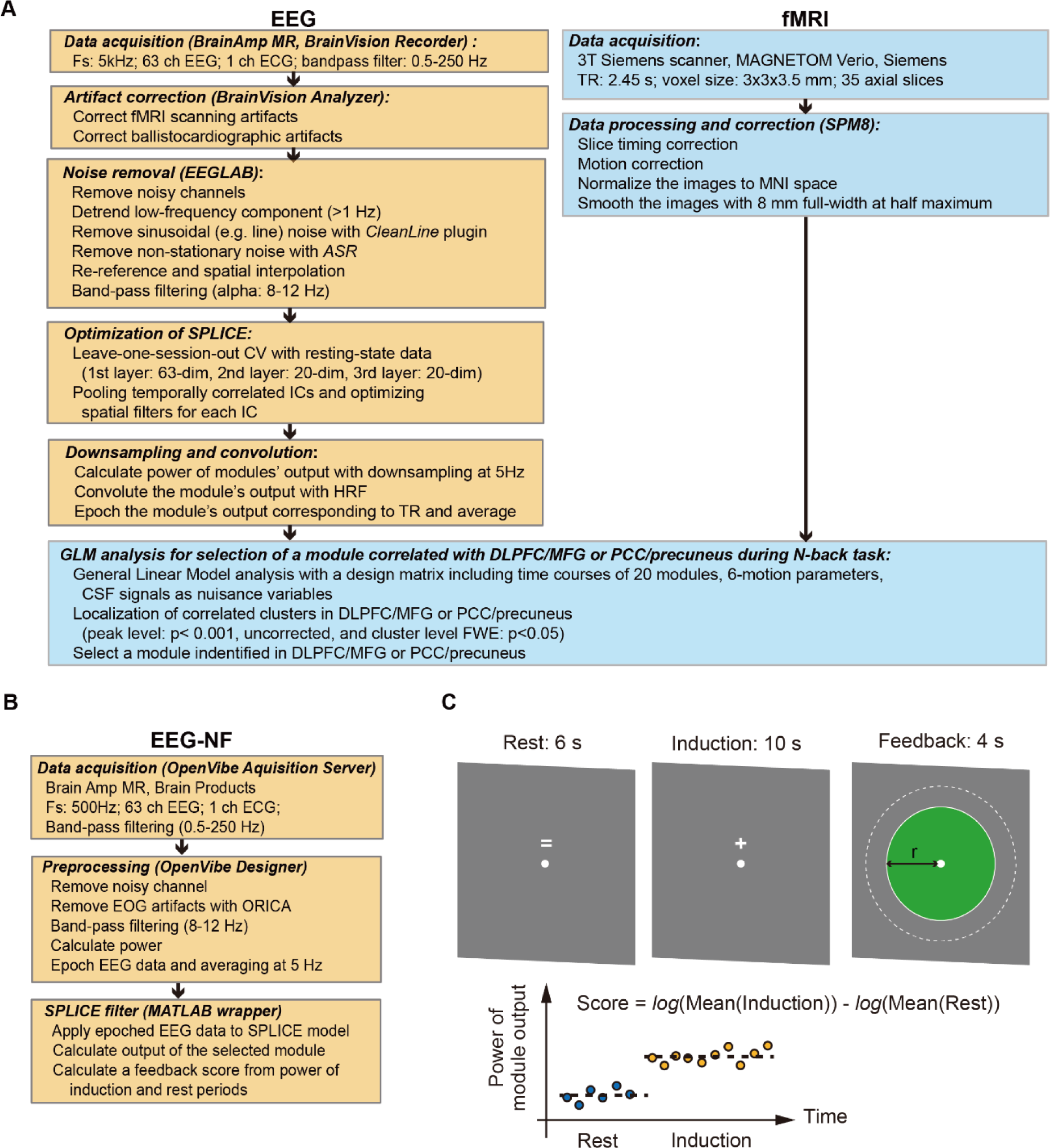
A flowchart of preprocessing of the electroencephalography-functional magnetic resonance imaging (EEG-fMRI) data and its integration analysis in this study. (**A**) A flowchart of preprocessing of the EEG and fMRI data for the optimization of the stacked pooling and linear components estimation (SPLICE). (**B**) A flowchart of online processing for the EEG neurofeedback (NF). (**C**) A diagram depicting the EEG NF paradigm. SPLICE, stacked pooling and linear components estimation; HRF, hemodynamic response function; MNI, Montreal Neurological Institute; FWHM, full width at half maximum; GLM, General Linear Model; BOLD, blood-oxygen-level-dependent; CSF, cerebrospinal fluid; FWE, family-wise error.

For each participant, we extracted the fMRI time courses within each region of interest (ROI). To remove several sources of spurious variance and their temporal derivatives, linear regression was performed, including six motion parameters in addition to averaged signals over gray matter, white matter, and cerebrospinal fluid. Furthermore, to reduce spurious changes caused by head motion, the data were using a method that reduces motion-related artifacts. A high-pass filter (< 0.008Hz) was applied to these sets of time courses prior to the regression procedure.

#### 2.4.2 Preprocessing of EEG data

To extract the EEG network features, the SPLICE model was optimized from the resting-state EEG data simultaneously recorded with fMRI, applying leave-one-session out cross-validation for each individual. Before the training of the SPLICE model, the raw EEG data were cleaned using Brain Analyzer software (Brain Products) to remove i) fMRI scanning artifacts and ii) ballistocardiographic artifacts. In the next step, we used the EEGLAB toolbox in MATLAB software (Delorme and Makeig, 2004) to process the data as follows: iii) remove noisy channels by visual inspection; iv) apply a high-pass filter (> 1 Hz) to detrend the EEG signal; v) remove harmonic noise with CleanLine (add-on in EEGLAB); vi) apply Artifact Subspace Reconstruction (ASR; Mullen et al., 2013) to remove stationary/non-stationary noise for continuous data; vii) interpolate removed channels to the original channel space and re-reference the data; and viii) apply a bandpass filter between 8 and 12 Hz (alpha band), as per a previous publication by de Munck et al. (2007).

#### 2.4.3 EEG-fMRI integration analysis for the selection of a module for NF training

We set 63, 20, and 20 dimensions for the first layer (sources), the second layer (modules), and the third layer (top-level ICs), respectively, similar to a previous study of resting-state brain networks using fMRI (Laird et al., 2011); we then estimated the parameters of the SPLICE model from the resting-state EEG data recorded in the fMRI scanning. We used a FCNef protocol (Yamada et al., 2017; Taylor et al., 2021) to create a functional localizer during the N-back task, and evaluated the SPLICE modules using EEG-fMRI data collected during the N-back task. We estimated the time courses of the output of modules through the SPLICE model and then downsampled them at 5 Hz. We applied the fMRI signal to the general linear model (GLM, SPM12) to identify a module that modulated the time course during the N-back task. The design matrix consisted of six motion parameters pertaining to head motions and the EEG network features, convoluted with the hemodynamic response function (HRF), and decimated to match the time instant of the fMRI signal (Mantini et al., 2007; Tsuchimoto et al., 2017). In the correlation analysis, we identified the modules that significantly correlated with brain activity, based on whether a statistical cluster existed in the DLPFC/MFG or PCC/Precuneus with a thresholding p-value < 0.001, uncorrected, and cluster-level p < 0.05 corrected by Family-Wised Error (FWE). Five participants with clusters identified in the DLPFC/MFG performed to upregulate the EEG network feature, and three participants with clusters identified in the PCC/precuneus performed to downregulate the one in the NF training. The participants were blinded to the group allocation and to their upregulation/downregulation status until the end of NF training.

Separately from this, we also validated the use of the SPLICE model in comparison to a standard ICA. Specifically, we calculated the mean log-likelihood of the SPLICE model, i.e., the negative of *L*(**x**) in Eq.(3) averaged over test data, using a leave-one-session-out cross validation scheme. Similar to this, we also obtained the mean log-likelihood of a standard ICA model, estimated by FastICA (the initial values of the first layer of the SPLICE model; Hyvärinen, 1999), with the same source prior as that in the top layer of SPLICE. For a fair comparison, we did not incorporate pooling across time points into the SPLICE model in this validation. We compared the cross-validated log-likelihood values, evaluated for all the individuals, between the SPLICE model and ICA using a paired t-test.

### 2.5 Neurofeedback protocol

Figure 3B shows the workflow of the online EEG NF training process. The real-time NF system consisted of OpenViBE software (v1.0.0; Renard et al., 2010) for real-time EEG signal acquisition, a feedback module programmed using MATLAB, and a Brain Amp MR (Brain Products) amplifier with a 63-ch EEG electrode cap. Raw EEG signals were acquired at 500 Hz using Brain Amp MR and the OpenViBE Acquisition Server. Using OpenViBE Designer (and interactive graphical user interface: GUI), the EEG data were subjected to electrooculography artifact removal (e.g., blinking, eye movement) using online recursive ICA (ORICA; Hsu et al., 2016) and a bandpass filter between 8 and 12 Hz. Subsequently, the signals were analyzed by MATLAB wrapper to predict EEG network features using the SPLICE model. According to a SPLICE module that was selected in advance, the L2-norm of the module was estimated as a targeting EEG network feature. It then displayed a feedback score depending on the task design. In this process, the output of the selected modules was downsampled at 5 Hz.

In the NF training, we modified the training protocols used in previous studies of FCNef (Yamada et al., 2017; Taylor et al., 2021) for EEG NF (see Figure 3C). We implicated an intermittent design as the NF training inducing reinforcement learning. We obtained a 5-min resting-state EEG recording and conducted five EEG NF sessions for all participants. We collected 20 trials in each session. A single trial consisted of “rest” for 6 s, “induction” for 10 s, and “feedback” for 4 s. Feedback was calculated as a score (-1 to 1) presented as a circle on a monitor. For the NF group, the score was a difference of the mean power of between “induction” and “rest” period with the logarithm scale in eleven steps (resolution = 0.2; if the score was less than -1 or more than 1, the circle size was minimum or maximum). For the sham group, the NF system randomly generated a score between -1 and 1. Participants were asked to try to increase the disc size as feedback on each trial by doing their best to “do something with their brain” during the induction period. In the instructions, participants were provided with examples such as performing mental calculations or recalling words to help imagine as to how to manipulate their own brain signals, at the beginning of the training. Participants did not receive specific, explicit strategies associated with the analysis of feedback scores during the NF training.

We randomly divided the participants into two training groups (NF and sham); however, we controlled the number of men and women in each group. Applying a two-sample t-test, we found no significant differences in depressive symptoms (BDI-II: p = 0.257; RRS: p = 0.317) between the two groups (Table 1). We compared scores from the psychological questionnaires to evaluate the effects of NT training on depressive symptoms using 2-way analysis of variance (ANOVA) for scores across time (pre/post) and groups (NF/sham); we used t-test (function of ‘anovan.m’, and ‘ttest2.m’ in the statistics toolbox of MATLAB) and effect size such as Cohen’s d for analyzing differences between pre-and post-scores (computeCohen_d.m; https://www.mathworks.com/matlabcentral/fileexchange/62957-computecohen_d-x1-x2-varargin). The sample sizes of the groups were small, so we assumed that it would be difficult to detect statistically significant differences using a typical statistical test, such as a t-test or ANOVA. To estimate a sufficient sample size, we calculated sample sizes from Cohen’s d in this comparison using G*Power 3.1.9.2 software (Faul et al., 2007; https://www.psychologie.hhu.de/arbeitsgruppen/allgemeine-psychologie-und-arbeitspsychologie/gpower.html). To evaluate participants’ performances in the N-back task, we calculated d’ (d-prime), a variable which is used in signal detection analysis (Palamedes toolbox in MATLAB; Prins and Kingdom, 2018).

**Table 1.** The demographics of the participants who underwent neurofeedback (NF) training

| Variables |  | NF group<br>(n = 9) | Sham group<br>(n = 9) | p-value<br>of t-test |
| --- | --- | --- | --- | --- |
| **Age (years)** |  | 28.62 (10.84) | 25.97 (7.22) | 0.606 |
| **Gender** | Women | 2 | 2 |  |
|  | Men | 7 | 7 |  |
| **BDI-II** |  | 7.89 (3.26) | 10.44 (6.50) | 0.257 |
| **RRS** |  | 40.67 (9.18) | 35.44 (12.91) | 0.317 |

NF: neurofeedback; BDI-II: Beck Depression Inventory II; RRS: Rumination Response Scale

**Table 2.** Comparison of statistical effect size associated with depressive symptom scores between the neurofeedback (NF) and sham groups

| Variables | NF group<br>(n = 8) |  | Sham group<br>(n = 9) |  | Two-way ANOVA<br>{time, group} | Two-sample<br>t-test<br>(post-pre) | Cohen's d |
| --- | --- | --- | --- | --- | --- | --- | --- |
|  | Pre | Post | Pre | Post | F-value (p-value) | t-score<br>(p-value) |  |
| **BDI-II (total)** | 5.25<br>(5.52) | 3.75<br>(4.43) | 7.22<br>(7.72) | 5.33<br>(6.34) | time: 0.70 (0.41)<br>group: 0.63 (0.43)<br>time x group: 0.01 (0.92) | 0.38<br>(0.710) | 0.184 |
| **-Cognitive depression** | 2.25<br>(3.37) | 1.75<br>(2.76) | 2.56<br>(3.47) | 2.44<br>(3.05) | time: 0.21 (0.65)<br>group: 0.08 (0.78)<br>time x group: 0.03 (0.86) | -0.80<br>(0.434) | -0.390 |
| **-Somatic depression** | 3.00<br>(3.46) | 2.00<br>(2.61) | 4.67<br>(4.66) | 2.89<br>(3.76) | time: 0.99 (0.33)<br>group: 1.17 (0.29)<br>time x group: 0.09 (0.76) | 0.99<br>(0.340) | 0.479 |
| **RRS (total)** | 31.00<br>(5.68) | 28.25<br>(4.40) | 31.33<br>(8.00) | 31.67<br>(8.70) | time: 0.60 (0.44)<br>group: 0.25 (0.62)<br>time x group: 0.41 (0.53) | -1.49<br>(0.158) | -0.722 |
| **-Brooding** | 7.50<br>(5.68) | 6.63<br>(1.92) | 7.78<br>(2.11) | 7.89<br>(2.93) | time: 0.94 (0.34)<br>group: 0.23 (0.63)<br>time x group: 0.39 (0.54) | -1.23<br>(0.220) | -0.626 |
| **-Depression** | 16.25<br>(2.96) | 15.88<br>(2.95) | 17.67<br>(5.48) | 17.56<br>(5.00) | time: 1.08 (0.31)<br>group: 0.03 (0.87)<br>time x group: 0.01 (0.93) | -0.21<br>(0.837) | -0.101 |
| **-Reflection** | 7.25<br>(1.98) | 5.75<br>(0.89) | 5.89<br>(1.45) | 6.22<br>(1.64) | time: 0.70 (0.41)<br>group: 1.21 (0.28)<br>time x group: 2.99 (0.09) | -2.16<br>(0.048) | -1.050 |

NF, neurofeedback; ANOVA, analysis of variance; BDI-II, Beck Depression Inventory II; RRS, Rumination Response Scale

## 3. Results

### 3.1 Comparison of the mean log-likelihoods with ICA and SPLICE

We compare the mean log-likelihoods of the SPLICE model to the standard ICA (FastICA in the first layer) with a leave-one-session-out cross validation using test data set across all participants. We found significant improvement (the paired t-test, t-score = 3.78, p = 0.0015; Supplementary Figure 7). We also obtained the mean log-likelihoods of initial values of the SPLICE model and found significant improvement (the paired t-test, t-score = 4.16, p = 6.5e-4). These results suggested that the SPLICE model sufficiently represents the resting-state EEG data simultaneously recorded with fMRI more than the conventional ICA.

### 3.2 Changes in feedback scores and psychological questionnaire scores

Figure 4 illustrates changes in depressive symptoms across time periods (pre- and post-NF) and groups (NF, sham) in the left panels, and differences in pre- and post-NF scores across the groups. Table2 summarizes statistical comparisons of BDI and RRS scores. We found an interaction among the RRS reflection scores (time × group, F-value = 2.99, p = 0.09) and a significant reduction (two-sample t-test, t-score = -2.16, p = 0.048) in these scores for the NF groups compared to the sham group (Figure 4G).

**Figure 4.**
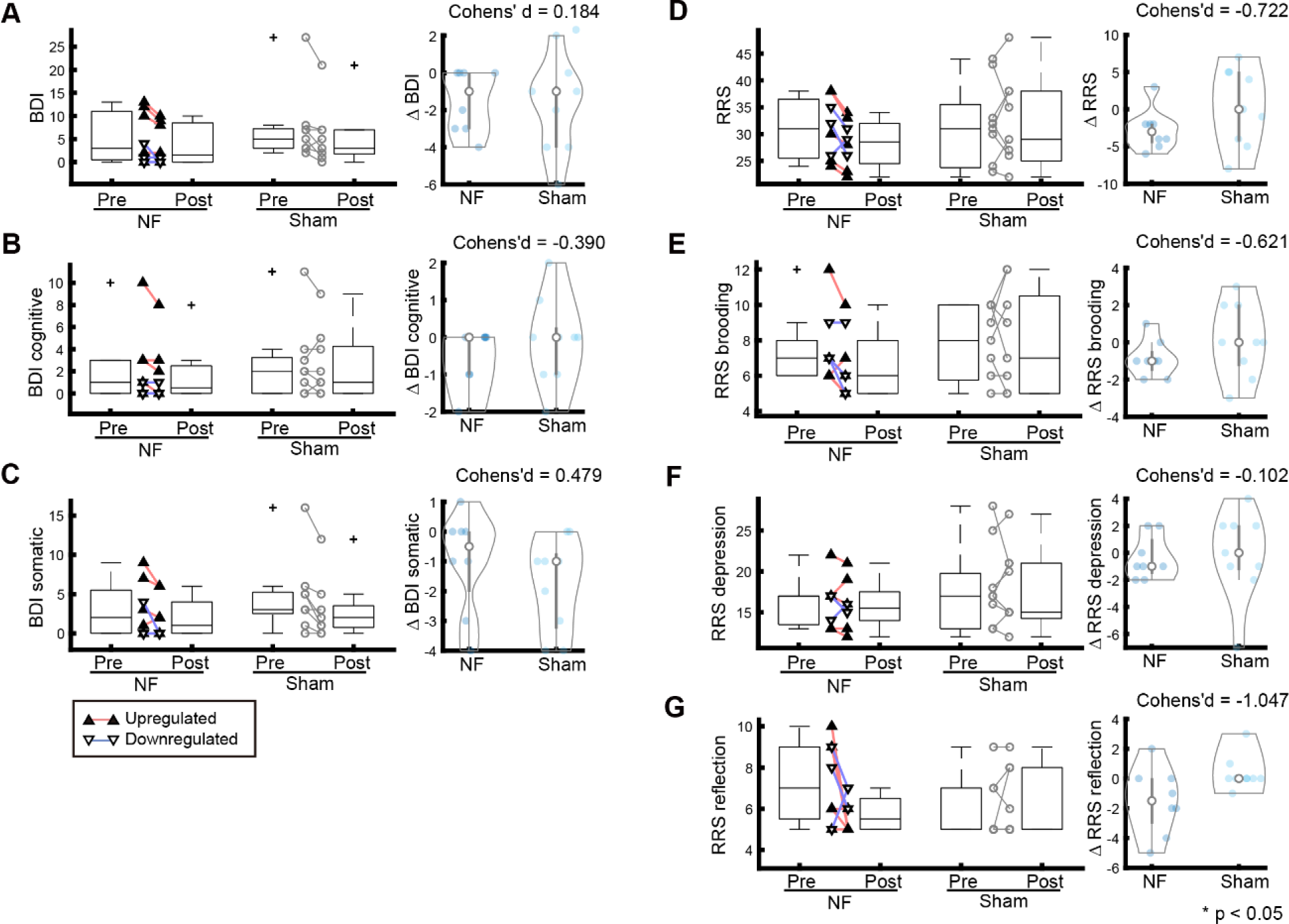
Effects of neurofeedback (NF) training on the depressive symptoms in the NF and sham groups. (**A**) Left panel shows boxplots and scatterplots displaying changes of in the Beck Depression Inventory II (BDI) score, and the right panel shows a violin plot displaying the differences between pre-and post-NF across the groups. (**B**-**G**) Same as (A) for BDI cognitive, BDI somatic, Rumination Response Scale (RRS), RRS brooding, RRS depression, and RRS reflection. In boxplots in the left panel in (**A**-**G**), the vertical line marks the median, 25th and 75th percentile, and cross presents outliers. In the violin plots in the right panel in (**A**-**G**), the compact boxplot shows median (circle), 25th and 75th percentile (bold lines), and data points (bright blue painted circle). An asterisk presents *p<0.05; two-sample t-test. BDI: Beck Depression Inventory II; RRS: Rumination Response Scale; NF: neurofeedback.

Two-way ANOVA results showed that NF training did not significantly affect the psychological questionnaire scores. We found large effect sizes in RRS scores (RRS: - 0.722, brooding: -0.626), but did not find sufficient effect sizes in the BDI scores (BDI: 0.184; cognitive depression: -0.390; somatic depression: 0.479). These results suggested that the RRS reflection scores in the group with NF training were reduced; however, these data do not imply an increase in the controllability of their EEG signals of participants during the NF training.

To identify changes in depressive symptoms depending on the type of regulation (upregulation/downregulation) of brain activity, we compared changes in the scores for each regulation type with changes in the scores in the sham group (Table 3). A large effect size and a significant difference were observed in the RRS reflection scores (Cohen’s d = -1.429, t-score = -2.56, p = 0.025) of the upregulation type. Similarly, a large effect size was found in the RRS scores (Cohen’s d = -0.765, t-score = -1.37, p = 0.195) of the upregulation type. However, we did not find a significant correlation between differences in the NF score and the RRS reflection score (Supplementary Figure 3G). Additionally, the other scores did not show a significant relationship with the NF scores. This suggests that our NF protocol may not have sufficiently effected depressive symptoms.

**Table 3.** Statistical comparison between the neurofeedback (NF) group (as per upregulation/downregulation) and the sham group.

| Variables | NF group<br>(upregulate,<br>n = 5) |  | Two-sample<br>t-test<br>(post-pre) | Cohen's d | NF group<br>(downregulate,<br>n = 3) |  | Two-sample<br>t-test<br>(post-pre) | Cohen's d |
| --- | --- | --- | --- | --- | --- | --- | --- | --- |
|  | Pre | Post | t-score<br>(p-value) |  | Pre | Post | t-score<br>(p-value) |  |
| **BDI-II (total)** | 7.40<br>(5.98) | 5.80<br>(4.49) | 0.24<br>(0.815) | 0.134 | 1.67<br>(2.08) | 0.33<br>(0.58) | 0.35<br>(0.736) | 0.232 |
| **-Cognitive depression** | 3.40<br>(3.91) | 2.60<br>(3.29) | -1.16<br>(0.270) | -0.645 | 0.33<br>(0.58) | 0.33<br>(0.58) | 0.16<br>(0.876) | 0.106 |
| **-Somatic depression** | 4.00<br>(3.87) | 3.20<br>(2.68) | 1.14<br>(0.276) | 0.636 | 1.33<br>(2.31) | 0.00<br>(0.00) | 0.38<br>(0.709) | 0.256 |
| **RRS (total)** | 31.00<br>(6.78) | 28.00<br>(5.52) | -1.37<br>(0.195) | -0.765 | 31.00<br>(4.58) | 28.67<br>(2.52) | -0.78<br>(0.455) | -0.518 |
| **-Brooding** | 7.40<br>(2.61) | 6.60<br>(2.07) | -0.95<br>(0.362) | -0.528 | 7.67<br>(1.16) | 6.67<br>(2.01) | -0.92<br>(0.380) | -0.613 |
| **-Depression** | 16.40<br>(3.72) | 16.20<br>(3.83) | -0.06<br>(0.954) | -0.033 | 16.00<br>(1.73) | 15.33<br>(0.58) | -0.27<br>(0.791) | -0.182 |
| **-Reflection** | 7.20<br>(2.17) | 5.20<br>(0.45) | -2.56<br>(0.025) | -1.429 | 7.33<br>(2.08) | 6.67<br>(0.58) | -1.04<br>(0.321) | -0.696 |

BDI: Beck Depression Inventory II; RRS: Rumination Response Scale; NF: neurofeedback

### 3.3 Relationships between feedback scores and depressive symptoms/behavioral performance

To clarify the effect of training on depressive symptoms, we examined the feedback scores for each group across days, as shown in Figure 5. At the end of the NF training, six of the eight participants showed slightly increased feedback scores compared to the scores from Day 1 and Day 3, but there was no significant difference. We did not find a significant improvement in the NF score within the NF group and also did not find noticeable differences in the feedback scores depending on the type of regulation of brain activity (up/down). Moreover, regarding correlations between differences (post – pre) in feedback scores and depressive symptoms, we examined the influence of the feedback scores on changes in symptoms. However, we did not find any significant differences (see Supplementary Figure 3) in the NF group. There were slight changes in NF scores (though not significant), but there were no changes in depressive symptoms. For this reason, the feedback scores did not reflect changes in depressive symptoms.

**Figure 5.**
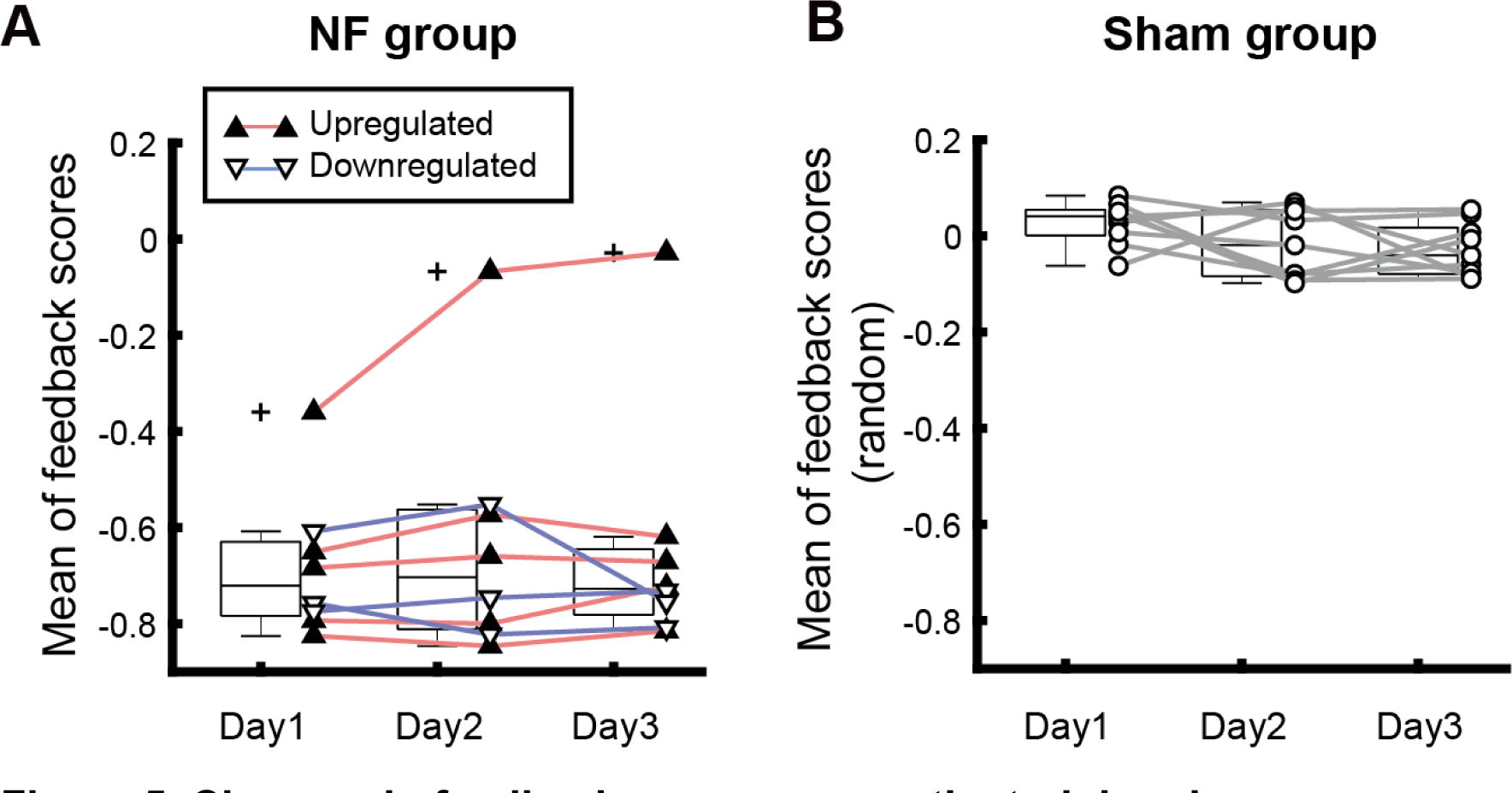
Changes in feedback scores across the training days. (A) Boxplots showing the distribution of the mean feedback scores across training days for the neurofeedback (NF) group. Red lines with black painted triangles and blue lines with white painted triangles represent each participant’s mean feedback score in a day. (B) Boxplots showing the distribution of the mean feedback scores across training days for the sham group. Gray lines with white painted circles show the mean feedback score in a day for each participant. The feedback scores were randomly generated; therefore, the mean scores are distributed around 0. Bars in the boxplots present the 25th percentile, median, and 75th percentile. NF: neurofeedback.

To evaluate changes in cognitive function, we evaluated changes in the discriminability index d’, which represents the behavioral performance in the N-back task (0-, 1-, 2-, and 3-back) at visit 4 (Day 1) and visit 7 (Day 4). We applied two-way ANOVA and two-sample t-tests across all conditions of the N-back task (see Supplementary Figure 4 and Supplementary Table 4). We expected to find a negative correlation between changes in feedback score and changes in d’ under conditions with higher cognitive load, such as 3-back. Differences in d’ for the 3-back task showed a large effect size (- 0.730) in the sham group. We found significant effects in the group across all conditions (0-back: 9.35, 1-back: 5.34, 2-back: 9.51, 3-back: 3.59; Supplementary Figure 5). These results suggest that the performance baseline for the N-back task in the sham group was significantly higher than that in the NF group. Therefore, we could not capture the effects of the NF training on the cognitive function in this experiment.

Among other scores not directly related to depressive symptoms, we found large effect sizes in AQ (total, attention switching) and OCI (total, obsessing) (Supplementary Table1). In the NF group, we observed other effects such as an increase in STAI2 trait (Cohen’s d = 0.612) compared to the sham group, but not in STAI2 state (Cohen’s d = 0.320). This may also be because the participants found it difficult to control their brain activity to increase feedback scores during the training.

## 4. Discussion

This study explored the application of the SPLICE model for EEG NF training, and investigated the effectiveness of this technique in depressive symptoms (particularly targeting rumination-related brain regions) in participants with subclinical depression. With the exception of one participant, none of the participants were able to control their training scores. In this section, we attempt to discuss the possible reasons behind the lack of success in controlling depressive symptoms using our NF paradigm in this study. We have categorized the section as follows: i) individual/environmental differences; ii) controllability of EEG signals during NF training; and iii) experimental procedure and the NF protocol design.

### 4.1 Individual/environmental differences affect SPLICE modeling

The analysis of the feedback scores suggested that the participants in the NF group were not able to control their own brain activity. We conducted SPLICE analysis in which the source signal was pooled and the temporally correlated networks corresponding to deactivation in the DMN or activation in the ECN observed in the fMRI signal at rest were extracted. Using GLM analysis, we selected a module that correlated with brain activity during the working memory task associated with modulation of the ECN. In this process, however, we realized that there were large individual differences among participants with reference to the modules and their correlated brain regions.

During the EEG-fMRI simultaneous recording (laying), the resting-state EEG data were used to optimize the SPLICE model, and EEG-fMRI data (laying) simultaneously recorded during the N-back task were used for module selection. Based on this, NF training was conducted by exclusively applying the EEG signal that was recorded outside the MRI scanner (sitting). Participants were asked to lie down inside the MRI scanner for the EEG-fMRI recording, which helped reduce artifacts caused by body/head movements. However, even slight movements by the participant caused significant artifacts in the EEG signal because of the high magnetic field. Therefore, the participants were asked to sit on a chair outside the scanner which caused less stress than being inside the scanner. Our data processing procedure did not sufficiently validate the robustness of the optimized model for different types of datasets such as tasked EEG inside/outside the scanner. Before every recording, as part of a quality check, we confirmed that the impedance of the EEG below was 10 kOhm. However, we did not sufficiently quantify the preprocessed EEG data. To resolve these problems, in future studies, the equivalence of the EEG signal quality should be evaluated by methos such as power spectrum analysis and ICA-based noise reduction (manually or machine-learning-based).

Another solution to minimize the data quality difference is the use of EEG/EEG-fMRI simultaneous NF (Wang et al., 2019; Zotev et al., 2020; Cury et al., 2020). For example, Wang et al. (2019) demonstrated proof of concept for the EEG NF protocol in patients with comorbid major depressive disorder (MDD) and anxiety symptoms; they compared three groups; frontal alpha asymmetry (ALAY group), high-beta downregulation training (Beta group), and control. Eighty-seven patients completed ten sessions of NF training twice a week, for five weeks. The ALAY and Beta groups showed a reduction of approximately 10 points in the BDI scores and a decrease of the Beck Anxiety Inventory (BAI) score by about eight points. Both psychological questionnaires showed significant interactions compared to the control group. This intensive and long-term NF training may be sufficient to induce significant effects on depressive symptoms. Zotev et al. (2020) continuously displayed fMRI signals in the left amygdala and left rostral anterior cingulate cortex (rACC), and frontal EEG asymmetries in the alpha and high- beta bands simultaneously as feedback for patients with MDD. A significant upregulation of the fMRI signal and frontal EEG asymmetries could enhance FC between the left amygdala and left rACC. Finally, they demonstrated a significant reduction in psychological measures such as the profile of mood state (POMS) and STAI (state), and correlations between EEG asymmetry changes and depressive symptoms. These NF trainings were performed in the same experimental environment and focused directly on the target regions. Further studies need to incorporate preprocessing ingenuity to reduce environmental differences, and to apply a sufficiently robust model.

### 4.2 NF design for NF training

Our NF protocol consisted of intermittent feedback with implicit strategies as detailed in fMRI NF studies based on reinforcement learning (Yamada et al., 2017; Taylor et al., 2021). In this framework, participants were required to concentrate on controlling the EEG signal for 10 s and then received feedback for 6 s. This paradigm is similar to that of a cognitive task such as working memory task. Therefore, we estimated the module from the tasked EEG data (n-back task); we assumed that our method could fit this framework such as reinforcement learning. On the other hand, a continuous feedback strategy seems be more suited for feedback-error learning, like motor control. In this case, short-time feedback with quick online processing is essential, and EEG a more suitable technique than fMRI (Zotev et al., 2020; Wang et al., 2019; Cury et al., 2020). This will allow participants to quickly understand how they need to control their brain time over time.

Additionally, previous studies have explicitly instructed participants on how to control brain activity during training. An advantage of the explicit strategy is that it makes it easy for the participants to imagine certain scenarios, such as positive autographical memory (Zotev et al., 2020; Wang et al., 2019), and motor imagery (Cury et al., 2020), and to monitor their own brain signals without a long delay. If the relationship between brain activity and its related cognition/image is firmly defined, it may help participants control their brain activity more easily during the NF. However, our protocol had a mismatch, and involved activation-based feature extraction and intermittent NF. We did not combine the advantages of both the above mentioned characteristics in the NF training. In future studies, we aim to resolve these methodological compatibility issues.

We did not find a significant improvement in the feedback score within the NF group with reference to the training effect (Figure 5). We examined the distributions of the feedback scores across three days for each participant (Supplementary Figure 6). We found that two participants did not exhibit positive scores during the NF training, five participants showed negative scores in most of the trials, and one participant showed scores spanning from the negative to positive. This suggests that we were not able to choose the proper module for each participant individually; therefore, most participants could not learn how to control their brain signals. In our study, only one participant showed improved feedback scores. If we could have chosen the modules precisely for each participant, all the participants would have been able to control their EEG signals day after day with improved efficiency.

Another possible reason for lack of success of the methodology is the selection of the frequency band. Based on previous studies (de Munck et al., 2007), we assumed that the alpha band of EEG signals correlated mainly with fMRI data. However, we did not find common changes in the time-frequency component, such as event-related desynchronization/event-related synchronization, across the participants during the NF training. Other frequency bands should also be examined in the optimization process to identify the best model for each participant.

### 4.3 Limitations of this study

A major limitation of this study is that we could not recruit a sufficient number of participants to evaluate our NF training protocol. We tested the NF training protocol in healthy participants with subclinical depression who showed BDI scores of more than three points, because our project aimed to focus on the maintenance of mental health in daily life of healthy individuals. We could not recruit participants with severe MDD in this study; therefore, the scores pertaining to depressive symptoms were lower than those in previous patient studies (Wang et al., 2019; Young et al., 2017; Zotev et al., 2020). Some participants had low scores from the beginning (Day 1) and did not show enough improvement due to the NF training, possibly because of the floor effect. One of the difficulties we faced was in recruiting appropriate participants; in actuality, we recruited and screened about a hundred potential participants for subclinical depression. Difficulties in recruiting the approximate population have been reported in previous studies as well; Wang et al. (2019) showed a drop-off rate of about 5% in MDD patients. To improve the sensitivity of pre-screening, the Self-rating Depression Scale (SDS; Zung, 1965) questionnaire should also be used in addition to the BDI questionnaire. We completed the NF training for 17 participants (a drop-off rate from the Screening to EEG-fMRI experiment: 27%, and that from the EEG-fMRI to the NF experiment: 64%). However, this reduction in sample size led to a reduction in statistical power during the evaluation. According to the effect size (such as Cohen’s d), we estimated sufficient sample sizes (p < 0.05, two-sample t-test) for each group to be as follows: BDI: 671 samples; BDI cognitive: 150 samples; BDI somatic: 100 samples; RRS: 45 samples; RRS brooding: 60 samples; RRS reflection: 22 samples. Thus, the sample number (NF: n = 8; sham: n = 9) was insufficient to support the evaluation of the effects of NF training with solid statistical power at a high significance level. In future studies, the procedure for participant recruitment should be improved.

From a systematic point of view, we considered noise reduction in the online process. We applied online recursive ICA (ORICA; Hsu et al., 2016) that optimized the weights of the independent components (ICs) with the real-time process and was practical to remove artifacts such as blinking or eye movements. To select artifact ICs, we focused on the time course of Fp1/Fp2, which was close to the eyes. With the time-window approach, we automatically selected the IC that was most correlated with the EEG signals of Fp1/Fp2. This method chose a proper IC when static EOG artifacts occur. However, it did not work if there was no EOG artifact in Fp1/Fp2. In an on-going study, we have improved EOG artifact removal.

We expected that functional regularization in the DMN or ECN might reduce depressive symptoms, such as those represented in the BDI and RRS. We found a large effect on RRS (total, brooding, and reflection) in the NF group compared to the sham group. Additionally, in the subgroup analysis (based on activity upregulation and downregulation), large effect sizes were observed in RRS total and reflection scores in the subgroup that showed upregulation of DLPFC/MFG. In a previous study, ROI-based fMRI NF training targeting the left DLPFC led to reductions in depressive symptoms, as reflected in BDI, the 17-item Hamilton rating scale for depression (HRSD), and RRS scores (Takamura et al., 2020). The authors applied fMRI NF for six MDD patients for five days. This training was aimed at upregulating an fMRI signal within the left DLPFC that was identified based on a functional localizer task such as the N-back task. They found a significant correlation between RRS rumination score changes and fMRI signal changes in the left DLPFC during the NF training. In this study, we performed EEG-based NF training without measuring the fMRI signal simultaneously; therefore, our results are not comparable to their results of this previous study. Examining the fMRI signal during the EEG NF training, may thus help identify the brain activity evoked by the training.

We attempted to develop an EEG NF method analogous to the fMRI NF method. In the integration analysis of EEG-fMRI data, HRF-convoluted frequency powers were calculated to match time instants between EEG and fMRI data, and then GLM was applied to identify brain regions or networks (Mantini et al., 2007). Another approach combines EEG-microstate analysis which categorized the data into four to six states based on the clustering algorithm, and then identifies brain regions using correlation analysis (Brechet et al., 2019; Al Zoubi et al., 2020). These methods may achieve dimension reductions and describe the relationship between fMRI and EEG data. In future studies, we plan to evaluate EEG-fMRI data in several categories pertaining to different conditions to quantify the accuracy of the SPLICE model.

In summary, we proposed a new framework of EEG NF training focusing on EEG network features combined with fMRI information such as the ECN and DMN. In the three-day NF training, we found a significant reduction in depressive symptoms as reflected by the RRS reflection scores; however, this reduction was not associated with the feedback score during the NF training. Because of suboptimal module selection for the NF training, the participants could not control their brain activity, and thus we did not see significant improvement in their depressive symptoms. In future studies, several points must be considered, such as changing of the NF protocol (intermittent/continuous), type of instruction (explicit/implicit), frequency bands of EEG, and artifact removal method.

## Author Contributions

Takeshi Ogawa and Hiroki Moriya designed the study. Takeshi Ogawa and Nobuo Hiroe performed data acquisition. Takeshi Ogawa and Jun-ichiro Hirayama analyzed the data. Takeshi Ogawa, Jun-ichiro Hirayama, and Motoaki Kawanabe performed data analysis and interpreted the results. Takeshi Ogawa and Jun-ichiro Hirayama wrote the original draft of this manuscript.

## Acknowledgments

We thank the following individuals for their contributions: Yoko Matsumoto for her help in recruiting participants, scheduling, and experimentation; Kana Inoue and Tachi Kaori for their help in screening participants during the Screening experiment; Mihoko Sato, Ayumi Yuki, and Yoshiko Itakura for their support in the preparation of the EEG-fMRI and NF training experiments; and Takashi Yamada for discussing the design the neurofeedback experiment. We would also like to thank the ATR Brain Activity Imaging Center (BAIC) team, and in particular, Akikazu Nishikido and Ichiro Fujimoto, for their technical support.

## Data Availability Statement

The data that support the findings of this study are available from the corresponding author, Takeshi Ogawa, upon reasonable request.

## Contribution to the Field Statement

Advanced treatments for depressive symptoms, such as real-time fMRI neurofeedback (NF), have been proven in several studies. Regularization of brain activity/functional connectivity (FC) between the executive control network (ECN) and the default mode network (DMN) during fMRI NF has been proposed to be effective in reducing depressive symptoms. However, it is difficult to install this system in practice because the cost is high, and no practical signal processing techniques have been developed to extract FC-related features from EEG. In this regard, stacked pooling and linear components estimation (SPLICE), recently proposed as a multilayer extension of independent component analysis and related independent subspace analysis, can be a promising alternative, although its application to resting-state EEG has never been investigated. This method may help modulate the target FC in EEG-based NF training. This study applied the SPLICE model in EEG NF training and explored its possible effectiveness in targeting depressive symptoms; rumination-related brain regions in participants with subclinical depression. We optimized the SPLICE model with simultaneous EEG-fMRI data to extract EEG network features such as a module correlated with fMRI signals in the DLPFC/MFG or in the PCC/precuneus by GLM analysis. As a result of the NF training using the extracted module, a large effect size and significant differences were found in the RRS reflection scores of the NF group.

However, no significant correlation was found between the training scores and depressive symptoms, suggesting that this observation may not be a training-specific effect. Other scores showed that RRS and RRS brooding scores had large effect sizes, but no statistically significant improvement was seen when the NF group was compared to the sham group. Therefore, our NF protocol was unsuccessful in improving depressive symptoms and cognitive function.

