## Supplementary information for "EEG-based neurofeedback with network components extraction: a data-driven approach by multilayer ICA extension and simultaneous EEG-fMRI measurements"

### *Supplementary Material*

**Supplementary Table 1. Statistical comparison of pre-and post-neurofeedback (NF) training questionnaire scores.**

| Variables | NF group (n = 8) | | Sham group (n = 9) | | Cohen's d |
|---|---|---|---|---|---|
| | Pre (Day1) | Post (Day4) | Pre (Day1) | Post (Day4) | |
| **AQ (total)** | 19.13 (6.81) | 17.25 (7.05) | 19.22 (10.53) | 19.00 (10.58) | -0.626 |
| **-Attention switching** | 4.63 (1.69) | 3.75 (1.58) | 4.22 (2.28) | 4.33 (1.66) | -1.113 |
| **OCI (total)** | 17.00 (14.61) | 13.5 (14.01) | 21.56 (14.36) | 23.67 (24.99) | -0.539 |
| **-Hoarding** | 2.00 (2.51) | 1.88 (2.95) | 3.22 (2.11) | 3.74 (0.44) | -0.209 |
| **-Obsessing** | 4.25 (4.10) | 2.65 (3.20) | 4.56 (4.64) | 6.69 (0.67) | -0.677 |
| **STAI2 (trait)** | 36.62 (4.27) | 38.13 (6.49) | 33.89 (9.71) | 32.78 (11.63) | 0.612 |
| **STAI2 (state)** | 43.75 (6.78) | 43.38 (5.55) | 40.56 (11.82) | 38.67 (14.08) | 0.320 |
| **BIS-11 (attention impulsive)** | 16.75 (2.38) | 15.50 (1.92) | 15.11 (2.62) | 15.11 (3.00) | -0.791 |
| **BIS-11 (Motor impulsive)** | 21.38 (2.45) | 21.63 (2.01) | 20.89 (4.46) | 22.78 (4.38) | -0.700 |
| **BIS-11 (non-planning impulsive)** | 26.25 (3.81) | 26.25 (2.82) | 24.44 (5.27) | 24.44 (4.64) | 0.000 |
| **ZTPI (future)** | 3.35 (0.52) | 3.27 (0.53) | 3.50 (0.57) | 3.59 (0.42) | -0.756 |
| **ZTPI (past negative)** | 2.86 (0.62) | 3.06 (0.57) | 2.69 (0.70) | 2.66 (0.87) | 0.764 |
| **ZTPI (past positive)** | 3.60 (0.41) | 3.63 (0.53) | 3.54 (0.60) | 3.64 (0.60) | -0.336 |
| **ZTPI (present fatalistic)** | 2.64 (0.50) | 2.72 (0.68) | 2.59 (0.45) | 2.54 (0.54) | 0.376 |
| **ZTPI (present hedonistic)** | 3.15 (0.37) | 3.31 (0.52) | 3.16 (0.52) | 3.22 (0.41) | 0.369 |

NF, neurofeedback; AQ, Autism-Spectrum Quotient; OCI, Obsessive-Compulsive Inventory; STAI2, State-Trait Anxiety Inventory; BIS-11, Barratt Impulsiveness Scale 11 (BIS-11); Zimbardo Time Perspective Inventory (ZTPI)

**Supplementary Table 2. Comparisons of Pre (visits 2 and 3)/Post (Day 4) functional magnetic resonance imaging (fMRI) data obtained during the N-back task.**

| Brain region | Peak coordinate (MNI) | | | Cluster size | | Details |
|---|---|---|---|---|---|---|
| | X | Y | Z | Voxels | Peak of T score (p-value) | |
| 2-back > 0-back | | | | | | |
| R Angular gyrus | 38 | -66 | 44 | 244 | 5.38<br>(4.03e-6) | R Angular<br>R Inferior parietal gyrus<br>R Supramarginal gyrus |
| | 46 | -46 | 44 | | 3.96<br>(2.13e-4) | R Inferior parietal lobule<br>R SupraMarginal gyrus<br>R Angular gyrus |
| | 46 | -40 | 36 | | 3.83<br>(3.04e-4) | R SupraMarginal gyrus<br>R Inferior parietal lobule<br>R Angular gyrus |
| R Middle frontal gyrus, 2 | 34 | 44 | 2 | 330 | 4.17<br>(1.49e-5) | R Middle frontal gyrus, 2<br>R Superior frontal gyrus, 2<br>R Middle frontal gyrus, 2 |
| | 26 | 62 | 4 | | 4.71<br>(2.60e-5) | R Superior frontal gyrus, 2<br>R Middle frontal gyrus, 2<br>R Medial superior frontal gyrus |
| | 30 | 50 | -2 | | 4.14<br>(1.32e-4) | R Middle frontal gyrus, 2<br>R Superior frontal gyrus, 2<br>R Inferior orbitofrontal, 2 |

**Supplementary Table 3. Comparison of N-back task performance during simultaneous electroencephalography-functional magnetic resonance imaging (EEG-fMRI) recording**

| | | NF group | | Sham group | | Cohen's d |
|---|---|---|---|---|---|---|
| Measure | Level | Visit 2 & 3 (pre) | Visit 7 (post) | Visit 2 & 3 (pre) | Visit 7 (post) | |
| d' | 0-back | 4.12 ± 0.1 | 4.32 ± 0.1 | 4.32 ± 0.1 | 4.28 ± 0.1 | 0.48 |
| | 1-back | 3.85 ± 0.6 | 4.11 ± 0.4 | 4.20 ± 0.4 | 4.28 ± 0.2 | 0.33 |
| | 2-back | 3.18 ± 0.9 | 3.67 ± 0.7 | 3.52 ± 0.7 | 3.81 ± 0.6 | 0.30 |
| p (hit) | 0-back | 0.93 ± 0.2 | 0.99 ± 0.0 | 0.99 ± 0.0 | 0.99 ± 0.0 | 0.50 |
| | 1-back | 0.90 ± 0.2 | 0.96 ± 0.1 | 0.98 ± 0.1 | 0.99 ± 0.0 | 0.36 |
| | 2-back | 0.81 ± 0.2 | 0.91 ± 0.1 | 0.88 ± 0.1 | 0.94 ± 0.1 | 0.22 |

**Supplementary Table 4. Comparison of pre-and post-neurofeedback (NF) N-back task performance during the behavioral experiment**

| **Variables** | **NF group (n = 8)** | | **Sham group (n = 9)** | | **Two-way ANOVA {time, group}** | **Two-sample t-test (post-pre)** | **Cohen's d** |
|---|---|---|---|---|---|---|---|
| d' | **Pre** | **Post** | **Pre** | **Post** | F-value (p-value) | t-score (p-value) | |
| **0-back** | 3.39 (1.35) | 3.11 (1.15) | 4.20 (0.69) | 4.42 (0.31) | time: 0.01 (0.91)<br>group: 9.35 (0.005)<br>time x group: 0.52 (0.48) | -0.78 (0.445) | -0.381 |
| **1-back** | 2.25 (3.37) | 1.75 (2.76) | 2.56 (3.47) | 2.44 (3.05) | time: 0.71 (0.40)<br>group: 5.34 (0.028)<br>time x group: 0.04 (0.838) | -0.80 (0.434) | -0.122 |
| **2-back** | 3.00 (3.46) | 2.00 (2.61) | 4.67 (4.66) | 2.89 (3.76) | time: 0.65 (0.425)<br>group: 9.51 (0.004)<br>time x group: 0.27 (0.607) | -1.502 (0.154) | - 0.271 |
| **3-back** | 31.00 (5.68) | 28.25 (4.40) | 31.33 (8.00) | 31.67 (8.70) | time: 0.39 (0.533)<br>group: 3.59 (0.066)<br>time x group: 0.81 (0.373) | -1.49 (0.158) | -0.730 |

**Supplementary Figure 1. Raster plots of Beck Depression Inventory (BDI) and Rumination Response Scale (RRS) scores at the Screening experiment**

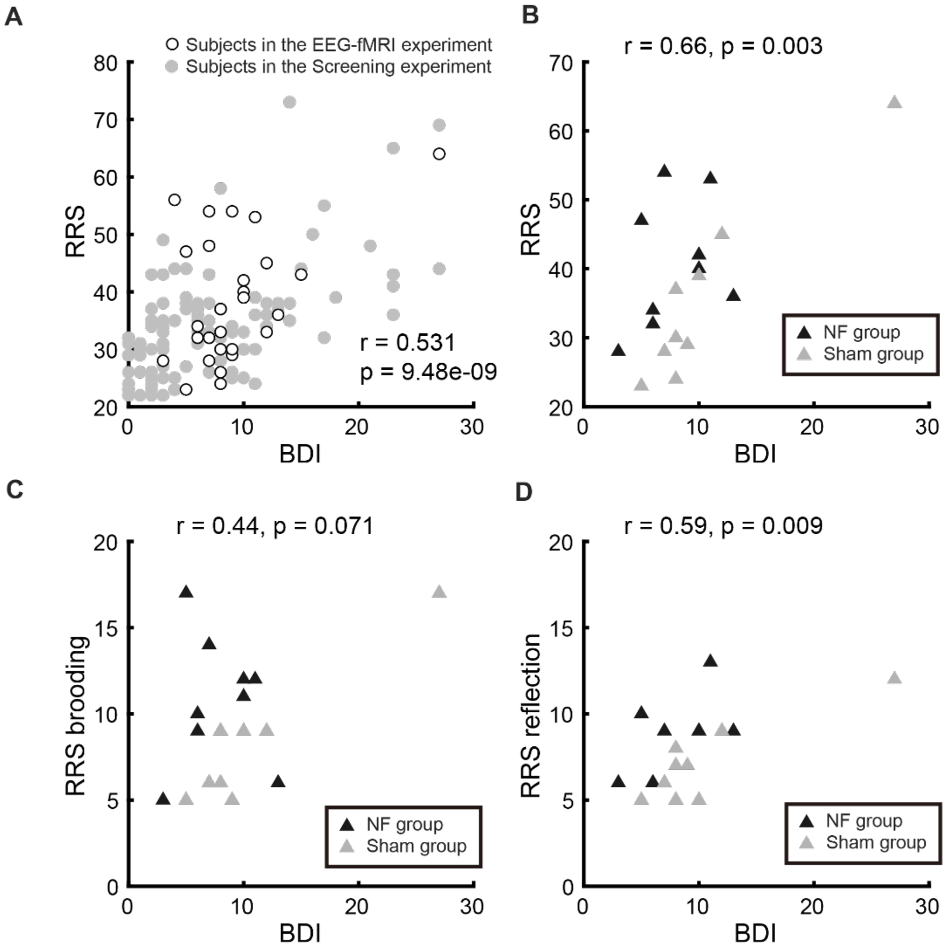


(**A**) Distribution of the BDI and RRS scores for participants in the Screening experiment. In the raster plots, white circles represent subjects who completed the EEG-fMRI experiment, and gray circles represent subjects who participated in the Screening experiment. Pearson's correlation between the BDI and RRS scores was calculated to evaluate the relationship between BDI and RRS. (**B**) A raster plot showing the BDI and RRS scores in the Screening experiment for participants who completed the NF experiment. (**C**-**D**) Same as (**A**) RRS brooding RRS reflection (black triangles: NF group, gray triangle: sham group).

BDI, Beck Depression Inventory II; RRS, Rumination Response Scale; NF, neurofeedback.

**Supplementary Figure 2. Identification of clusters illustrated the interaction of groups and times.**

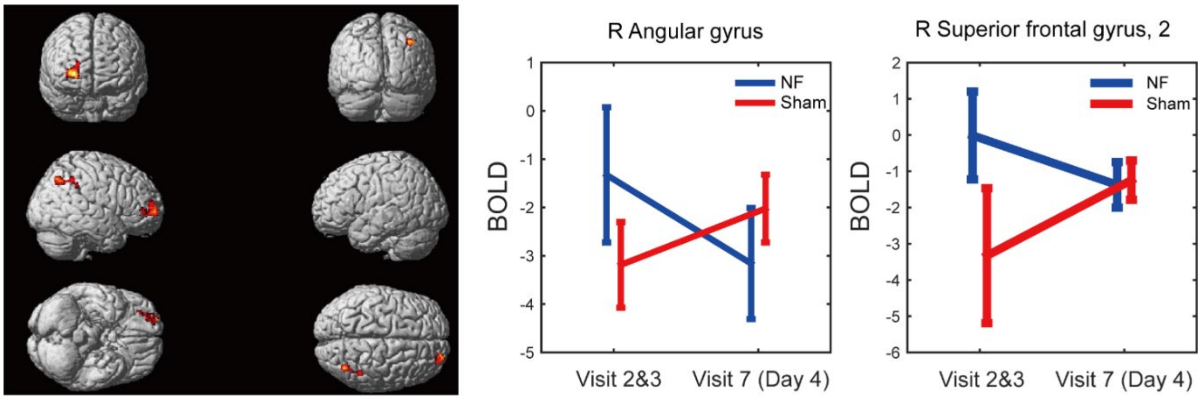


Clusters showed a significant interaction of group (neurofeedback: NF, sham) and time (pre: visits 2 and 3, post: visit 7, Day4). The contrast was 2-back > 0-back and labeled by Anatomical Automatic Labeling (AAL) toolbox; thresholding by p-value < 0.001, uncorrected and cluster level p < 0.05 corrected by Family-Wise Error (FWE, cluster size > 244 voxels). The left panel shows clusters that had interactions based on time and group. The middle and right panels show the BOLD signal of this contrast for each time and group. We did not collect the electroencephalography-functional magnetic resonance imaging (EEG-fMRI) data on visit 4 (Day1) for practical reasons (scheduling of experiment and less burden on participants). To compare the fMRI data on visits 2 and 3 as pre-NF training and visit 7 as post-NF training in this analysis, we statistically evaluated the interactions of groups (NF, sham) and time (visits 2 and 3, visit 7) and applied a full factorial analysis for fMRI data during the N-back task using SPM 12. Two clusters with a contrast of 2-back > 0-back were identified at the right angular gyrus (middle panel) and superior frontal gyrus 2 (right panel); labeled by AAL toolbox; thresholding by p-value < 0.001, uncorrected and cluster level p < 0.05 corrected by FWE (cluster size > 244 voxels; Supplementary Table 2. Regarding the behavioral performance in the N-back task during the EEG-fMRI simultaneous recording, we did not find significant effects in d’ or hit rates due to the NF training because of the ceiling effect (Supplementary Table 2 and 3). It is suggested that the neural basis of high cognitive load (2-back) could be processed with less effort in these regions due to the NF training.

**Supplementary Figure 3. Relationships between differences in depressive symptoms and changes in feedback scores.**

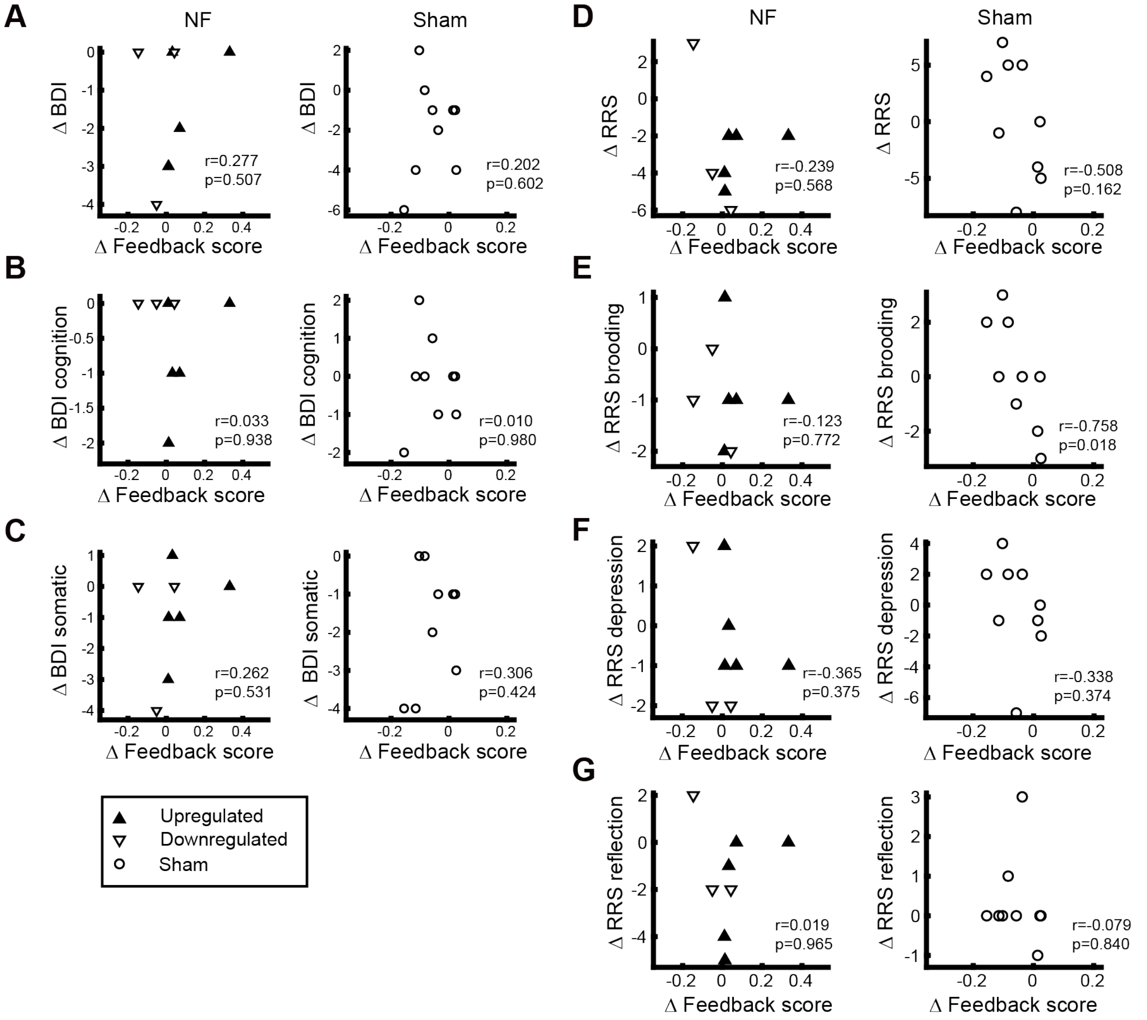


(**A**) The left panel illustrates a raster plot presenting a relationship between differences (post – pre) in feedback score and Beck Depression Inventory-II (BDI) for the neurofeedback (NF) group (black triangle: upregulated feedback; white triangle: downregulated feedback), and the right panel illustrates the relationship for the sham group. (**B-G**) Same as (**A**) for BDI cognitive, BDI somatic, Rumination Response Scale (RRS), RRS brooding, RRS depression, and RRS reflection. Pearson's correlations (r) and the corresponding p-values (p) are described at the bottom right of the plots.

**Supplementary Figure 4. Relationships between changes in task performances during the N-back tasks and changes in feedback scores.**

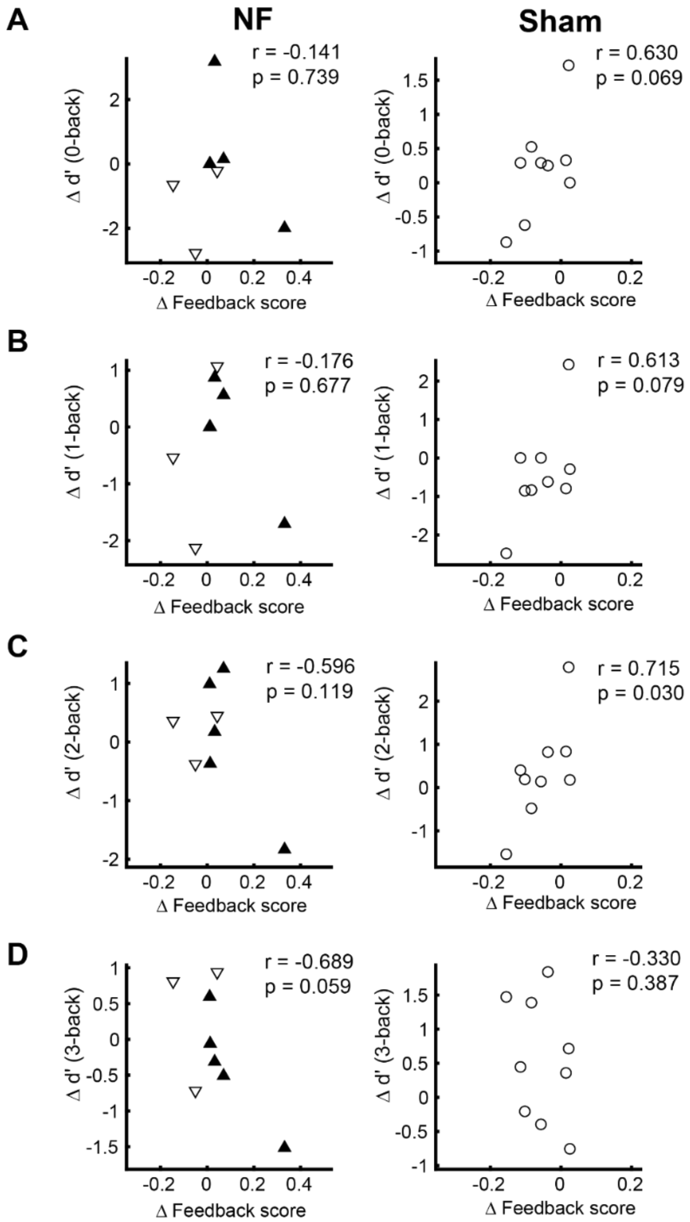


(**A**) The left panel illustrates a raster plot depicting the relationship between differences in feedback scores (Day 3 – Day 1) and d' of the 0-back task as a behavioral task for the neurofeedback (NF) group (Day 4 – Day 1) (black triangle: upregulated feedback; white triangle: downregulated feedback), and the right panel illustrates the relationship for the sham group. (**B**-**D**) Same as (**A**) for the 1-back, 2-back, and 3-back conditions. Pearson's correlations (r) are described at the bottom right of the plots.

**Supplementary Figure 5. Performances of participants with reference to the N-back task in the behavioral experiment at pre-and post-neurofeedback (NF) training.**

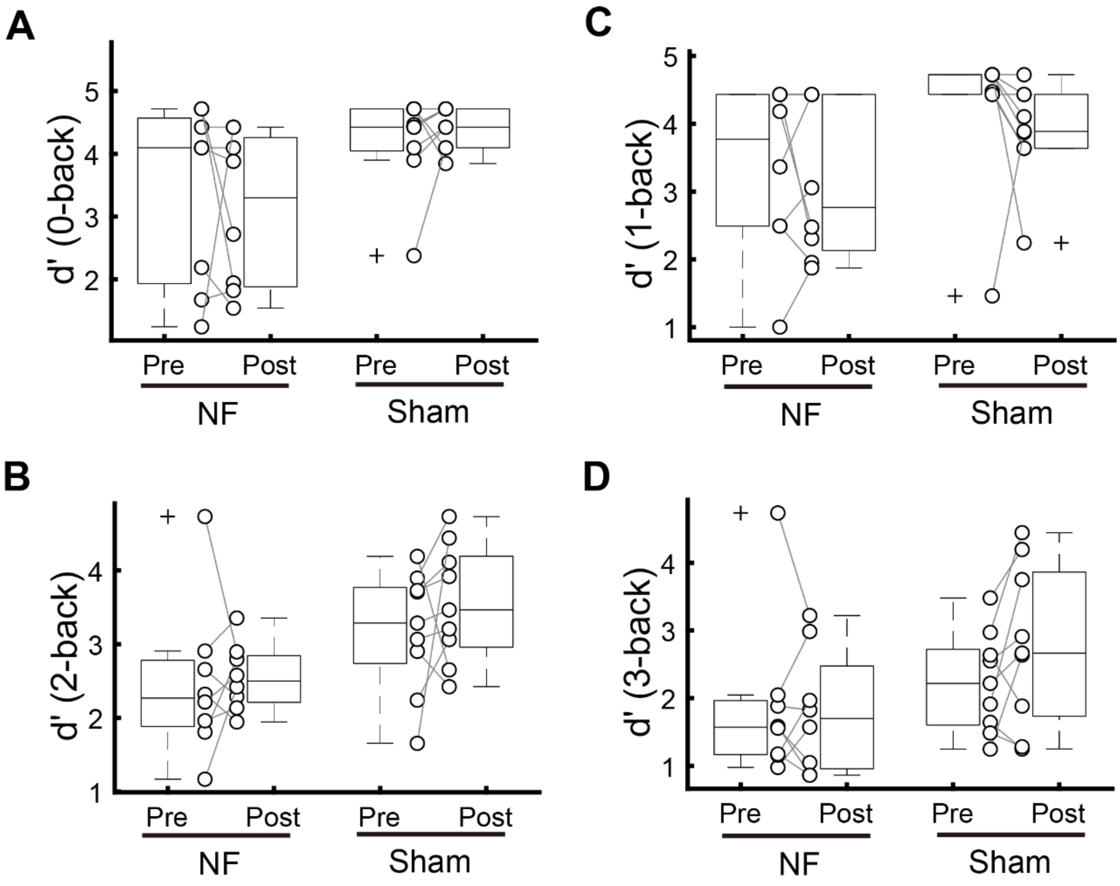


(**A**) Boxplots illustrating the distribution of d’ before and after neurofeedback (NF in the NF group and pre-and post-NF in the sham group during the 0-back task during the behavioral experiment (outside of the magnetic resonance imaging (MRI) scanner). Plots correspond to d’ for each participant. (**B**-**D**) Same as (**A**) for the 1-back, 2-back, and 3-back tasks.

**Supplementary Figure 6. Distributions of feedback scores across the neurofeedback (NF) training days**

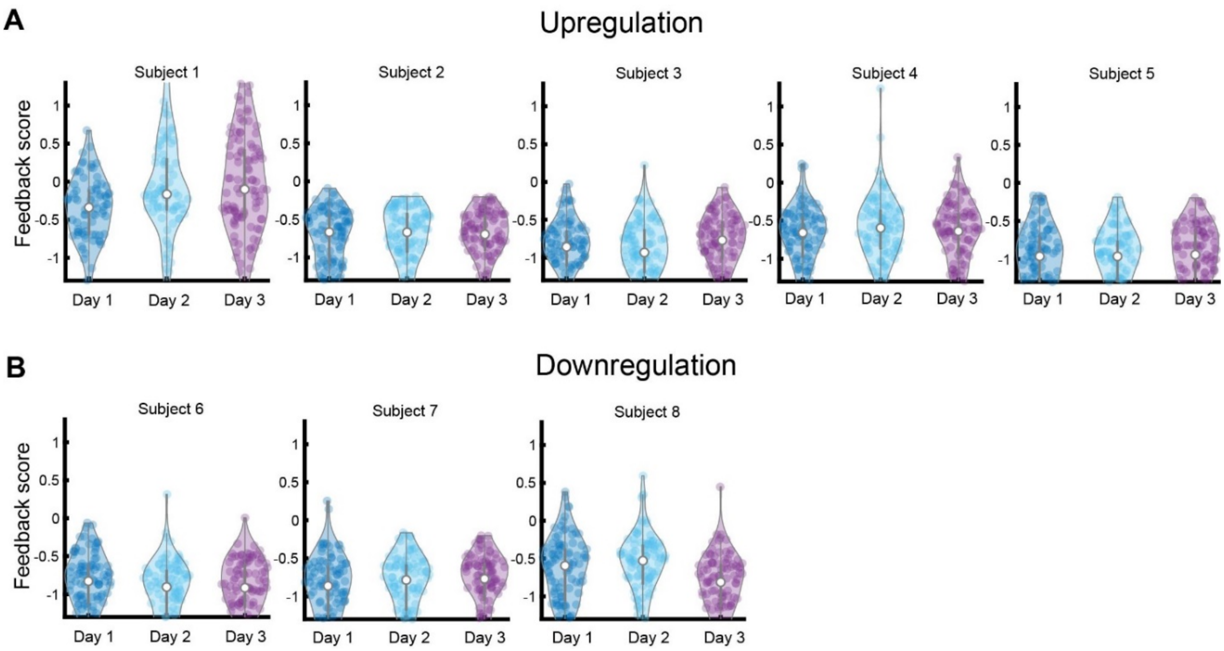


(**A**) Violin plots illustrating distribution of feedback scores across days for each participant in the upregulation subgroup, in the neurofeedback (NF) group. (**B**) Same as (**A**), violin plots of feedback score distribution for the downregulation subgroup in the NF group.

**Supplementary Figure 7. Distributions of the mean of log-likelihoods of the standard independent component analysis (FastICA, ICA with the maximum likelihood estimation) and the SPLICE model for the test data set.**

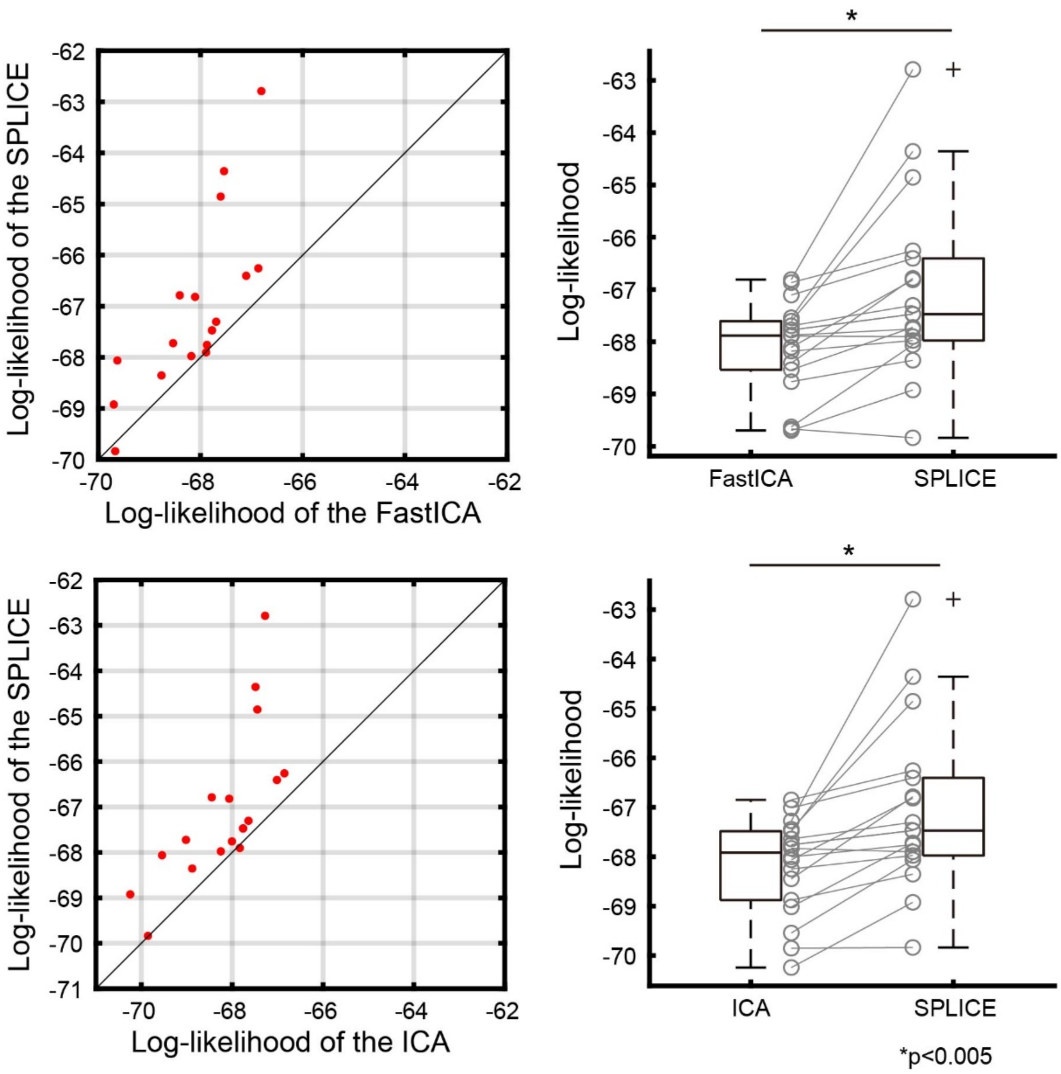


Raster plots in the left panel show distributions of the mean of log-likelihoods of the SPLICE model (y-axis) vs. the first layer ICA (FastICA, x-axis) for the test dataset in an upper panel, and the SPLICE model (y-axis) vs. a standard ICA (initial values estimated by FastICA, and then fine-tuned by the maximum likelihood estimation (MLE) as well as the SPLICE, with only the 1st ICA layer and the same top-level source prior, x-axis) in a bottom panel, across all participants in the NF training. Models painted in the upper left area over the black line represent better to explain EEG data. In the right panels: a boxplot in the upper right panel shows differences in the mean of the log-likelihoods between the FastICA and SPLICE model across the participants. Same as this, a boxplot in the bottom right panel shows differences in the mean of the log-likelihoods between the ICA (estimated by the MLE) and SPLICE model across the participants. Gray cycles correspond to each participant (*$p < 0.005$ with a paired t-test).
